# Symbiont-conferred immunity interacts with the effects of parasitoid genotype and intraguild predation to shape pea aphid immunity in a clone-specific fashion

**DOI:** 10.1101/2021.05.01.442267

**Authors:** Samuel Alexander Purkiss, Mouhammad Shadi Khudr, Oscar Enrique Aguinaga, Reinmar Hager

## Abstract

Host-parasite interactions represent complex co-evolving systems in which genetic variation within a species can significantly affect selective pressure on traits in the other (for example via inter-species indirect genetic effects). While often viewed as a two-species interaction between host and parasite species, some systems are more complex due to the involvement of symbionts in the host that influence its immunity, enemies of the host, and the parasite through intraguild predation. However, it remains unclear what the joint effects of intraguild predation, defensive endosymbiosis, within-species genetic variation and indirect genetic effects on host immunity are. We have addressed this question in an important agricultural pest system, the pea aphid *Acyrthosiphon pisum*, which shows significant intraspecific variability in immunity to the parasitoid wasp *Aphidius ervi* due to immunity conferring endosymbiotic bacteria. In a complex experiment involving a quantitative genetic design of the parasitoid, two ecologically different aphid lineages and the aphid lion *Chrysoperla carnea* as an intraguild predator, we demonstrate that aphid immunity is affected by intraspecific genetic variation in the parasitoid and the aphid, as well as by associated differences in the defensive endosymbiont communities. Using 16s rRNA sequencing, we identified secondary symbionts that differed between the lineages. We further show that aphid lineages differ in their altruistic behaviour once parasitised whereby infested aphids move away from the clonal colony to facilitate predation. The outcome of these complex between-species interactions not only shape important host-parasite systems but have also implications for understanding the evolution of multitrophic interactions, and aphid biocontrol.

## 1. Introduction

Natural ecosystem dynamics and their evolution are driven by complex interactions of selective pressures on interacting species caused by both environmental and within-species genetic variation e.g. [1]. A textbook example extensively investigated at a theoretical and empirical level is the interaction between hosts and parasites. Here, the fitness of a parasite is dependent on its host and, despite a vast range of evolved anti-parasite responses; organisms continue to be successfully parasitised [2]. Parasites often manipulate their hosts to improve their fitness [3] through *e.g*. promoting predator avoidance responses in the host, increasing its likelihood of survival [4]. Particularly complex interactions often occur in parasitoidism, a process that represents aspects of parasitism and predation. Complete parasitoidism occurs when the larva of the parasitoid develops within its parasitised host as a parasite. This results eventually in host death through mummification, where only the exoskeleton remains after parasitoid emergence [5,6]. An important example of a parasitoid is the hymenopteran endoparasitoid *Aphidius ervi* (Haliday), which is widely used as a biological control agent of the pea aphid *Acyrthosiphon pisum* (Harris) [7], in which it generally lays only one egg [6].

The co-evolutionary dynamics between antagonist species, such as an aphid host and its parasitoid, may drive evolution through a process of reciprocal adaptation and counter-adaptation, with selection for the development of resistance in the host and virulence traits in the parasitoid [8,9]. While resistance is widespread among host species, with evidence for degrees of endogenous resistance in some insect species, resistance is more often conferred or mediated by specific microbial symbionts [10,11]. Pea aphids show considerable within-species variability in resistance to the parasitoid *A. ervi* [12,13], and it has recently been shown that this variation is often explained by different protective symbionts found in different aphid lineages [11,13,14]. Therefore, the natural enemy of the host is an enemy of the symbiont. Lineages of the pea aphid exist in clonally reproducing populations under temperate favourable conditions, and these clones carry vertically transmitted (secondary) facultative symbionts in addition to its (primary) obligate symbiont *Buchnera aphidicola* [15,16]. The primary symbiont provides the aphid with nutrients that are lacking in its diet and that it could not otherwise produce [15]. Secondary symbionts have been implicated in various functions of aphid biology, including aiding in host-plant specialisation and particularly resistance to parasitoids [11,15,17].

Within-species genetic variability is known to affect both focal and other interacting species and communities [18, 19] and can be highly influential in determining the dynamics of host-parasitoid systems [20–22]. Further, indirect genetic effects theory outlines how the genotype of an individual can influence the phenotype of another individual (e.g. [23,24]), of the same species (indirect genetic effect, IGE) or another species (indirect interspecies genetic effect, IIGE). In the aphid-parasitoid system, the parasitoid wasp *A. ervi* alters aphid behaviour by influencing where aphids go to die during wasp larval development [21]. Indeed, such a behavioural modification of the aphid is influenced by the genotype of the wasp [21] and thus represents an IIGE [21,25,26]. Therefore, we need to consider the effects of both within-species genetic variability and their indirect effects when studying complex host-parasite systems.

Following the Red Queen hypothesis, interacting species must constantly evolve to maintain their position as a form of an evolutionary arms race between the species [27], which may result in either reciprocal selective sweeps [28] or sustained genotype oscillations [29,30]. In a host-parasitoid system, there is an arms race between host resistance (ability to survive the attack by the parasitoid) and parasitoid virulence (infectivity; the ability to overcome host defences) [27]. Most experimental examples of such co-evolutionary arms races [31] have demonstrated these as pairwise interactions between the parasitoid and its host. However, in natural ecosystems, parasitoids do not only operate in such pair-wise interactions as they are themselves interacting with other species that, for example, share aphid populations as prey [32,33]. Therefore, we predict that host and parasite fitness will be significantly affected by the presence of a common enemy such as the aphid lion larva *Chrysoperla carnea* (Stephens). Aphid lions naturally have an advantage over parasitoid wasps. The aphid lion can consume both healthy and parasitised aphids (parasitoid puparia), an ecological process known as intraguild predation (IGP) [33]. This leads to reducing the availability of viable, healthy aphid hosts for the parasitoid and, concomitantly, indirect reduction of parasitoid fitness [33,34]. In response to parasitoidism, aphids are known to exhibit altruistic risk-taking behaviour by exposure to predators when parasitised, thus, in essence, sacrificing the diseased few for the benefit of the genetically identical population (clone) [35,36].

Further, the bacterial symbiont, conferring a degree of immunity on the aphid host against its parasitoid enemy, adds another layer of complexity to the eco-evolutionary dynamics of the species interactions when intraguild predation occurs as the aphid lion is an enemy to the aphid host, the endosymbiont communities and the parasitoid. The complex interaction effects between within-species genetic variation of host and parasitoid when intraguild predation occurs may lead to ‘guild or diffuse co-evolution’ rather than pairwise co-evolution between two species [37,38,39], and may have important evolutionary implications for the pressures shaping aphid phenotype evolution in multi-trophic systems. To date, no experimental studies of such complex systems exist.

To address this gap in knowledge, we established a population of parasitoid *A. ervi* daughters using a half-sib quantitative genetic design, *sensu* Khudr *et al*. 2013 [21], and exposed two ecologically distinct lineages of the pea aphid *A. pisum*, with different defensive endosymbiont communities, to the effects of parasitoid intraspecific genetic variability to study host immunity with and without the presence of an intraguild predator (the aphid lion *C. carnea*). We hypothesised that, subject to the intraguild predator, differences in defensive endosymbiont communities and differences in parasitoid genotype may differentially affect pea aphid reproductive success and behaviour in a lineage-specific manner.

## 2. Materials and Methods

### Study organisms

#### Pea aphids and defensive endosymbiont

Two clonal lineages of pea aphid were selected for the experiment, N116 and our Q1 isolate. The N116 aphid is of the biotype (K) as it was originally isolated from alfalfa *Medicago sativa* (L.) by Dr Julia Ferrari in Berkshire UK [40]. It has been a laboratory lineage for *ca*. 10 years and was provided to us by Dr Colin Turnbull of Imperial College London. Q1 is of the biotype (G) [41], which was established from one female of a population colonising pea plants (*Pisum sativum* L.) isolated from the quadrangle garden of the Faculty of Biology, Medicine and Health, University of Manchester. The aphids were reared on faba bean *Vicia faba* var minor (Harz) obtained from a local supplier, Manchester, UK, and maintained at 22-24°C with a photoperiod of 16h (light) : 8h (dark). These two lineages of pea aphid are ecologically different. N116 is reported to have the heritable defensive endosymbiont *Hamiltonella defensa* [40,42] that confers relative immunity to parasitoidism. By contrast, in a pilot study, we established that Q1 was highly susceptible to being parasitoidised. Under temperate mesic conditions, aphids reproduce through parthenogenesis resulting in populations of genetically identical individuals.

#### Parasitoid wasp *A. ervi*

We purchased 250 mummies of aphids harbouring *A. ervi* developing juveniles from Koppert UK Ltd. Unlike the non-parasitic males, the females of this solitary koinobiont parasitoid wasp are an efficient natural enemy and biocontrol agent of pea aphids [33,43]. The female oviposits one egg in the viable aphid host. Subsequently, a larva hatches and parasitises the host consuming it internally whilst the parasitoid juvenile pupates, then develops into an adult that ecloses from the dead body of the host to resume the life cycle. Immediately upon their arrival, we separated the mummies into multiple 90mm petri dishes, each dish containing a small ball of dental cotton, approximately 20mm in diameter, which was saturated in 10% sucrose solution. The petri dishes were kept in the fridge at 10°C to slow the rate of eclosion from the aphid mummy (*i.e*. the wasp puparium). The petri dishes were taken from the fridge hourly and checked for the eclosion of wasps; the gender of the emergent wasp was observed; if all the individuals were of the same sex, then they could be used in the next stage of the experiment. The females were always isolated and kept separately from the males to insure the females were virgins prior to mating according to the quantitative genetic design explained below.

#### Intraguild predator *C. carnea* larva

The intraguild predator in our experiments was the aphid lion larva. The larvae were purchased from Ladybird Plant Care (UK) in tubes of approximately 300-500 individuals. The tube was emptied into a plastic container that contained some plant shoot parts with aphids as a provision and then kept in the fridge at 5°C until they were needed; this was to slow the rate of metabolism and prevent the larvae from cannibalising each other. The larvae were used within 48 hours of delivery or they were disposed of. As the wasps take ~11 days to emerge from the mummies, the aphid lion larvae (1^st^ instars) were ordered so that they would arrive on day 10 ready to be timely used where applicable in the experiment as described below.

### Experimental Design

Haplodiploidy is the sex-determination system in the Hymenopteran parasitoid wasp *A. ervi*, meaning that males are the result of unfertilised eggs and hence haploid (1n), while females are diploid (2n) since they produced from fertilised eggs [44]. Based on Khudr et al. (2013) [21], we mated randomly selected male wasps (sires) with randomly selected female wasps (dams) to establish a quantitative genetic half-sibling design. Each of the 34 sires was mated with a minimum of three dams, dependent on wasp availability right after their eclosion. We thus established sire-dam groups. Before the wasps were mated, they were isolated into Eppendorf tubes and inspected using a magnifying glass to observe abdomens and determine their sex; the female’s abdomen ends with a pronounced point (ovipositor) while the male’s abdomen is more rounded. The wasps were then put into the same tube by opening both tubes and putting them end to end. Once both wasps (sire and dam) had moved into the same tube, it was sealed with a small piece of foam. The mating wasps were monitored carefully until they completed copulation to ensure the dams were inseminated by the corresponding sire. Copulation was checked to have occurred within two hours of eclosion. If copulation did not happen, the female wasps were disposed of because of the short window of time during which the otherwise arrhenotokous parthenogenetic female wasp will be usually receptive to mating [21,44]. Once copulation was complete the foam was removed, the tubes were placed end to end, and we waited for the wasps to enter separate tubes before closing the lids and labelling the sire with its unique number (S1 – S*n*), and the dams with the number of the associated sire they mated with plus their own unique number in order of mating (*e.g*. S1 D1 – S*n* D*m*). Electronic supplementary material, figure S1 illustrates the experimental design.

Once mated, the inseminated dams were placed in their respective microcosms. The microcosms were constructed by removing the ends of a 2-litre PVC bottle and attaching one end to the plant pot and covering the other with a fine nylon mesh (‘Non-Fray’, Insectopia, UK). Each microcosm contained a 3-week-old broad bean plant that had been infested with 30 third instars of N116 just before putting the wasp into the enclosure. To release the dam into the microcosm the top section was held in place over the plant (leaving a small gap on one side), the lid of the tube was opened and sealed with the end of a finger and then the tube was passed through the gap onto the soil. Once the inseminated wasp was inside the microcosm, the top section of the microcosm was secured to the plant pot using 48mm wide polypropylene tape. The microcosms were placed, evenly spaced, into large trays, containing a shallow layer of water, in the growth chamber for eleven days. The conditions in the chamber were 22-24°C with a 16h (light):8h (dark) photoperiod; the water level in the trays was maintained and the positions of the microcosms on the trays were randomised every other day. On the eleventh day, the microcosms were taken from the growth chamber, opened, and all the mummies present were removed from the plant and inner surfaces of the microcosm using a fine damp paintbrush. Each mummy was placed in a separate 35mm petri dish that contained a small ball of dental cotton (approximately 10mm in diameter) saturated with 10% sucrose solution and labelled with the associative sire-dam number. The petri dishes were left at room temperature on the lab bench and left until we observed eclosion. Once the progenies (sib and half-sib daughters denoting the intraspecific genetic variability of the parasitoid) had emerged from the aphid mummies, they were individually introduced into a microcosm with a 3-week-old faba bean plant that had been infested with 30 third instars of N116. The microcosms were sealed, and each of the introduced daughters (i.e. parasitoid genotype) was given 11 days to parasitise the provided aphid population leading to the production of mummies. We then censused the aphid population per microcosm (mummified and healthy) and recorded the positions of the mummies off plant versus on plant. The whole procedure was repeated for the Q1 lineage. As such, the wasp daughters (parasitoid genotype) represented the intraspecific genetic variation effects in the parasitoid wasp, whereas the within-species genetic variation in the pea aphid host was presented by the inclusion of the N116 and Q1 lineages.

The remainder of the generated parasitoid daughters were used to test the effect of the presence of the aphid lion as an intraguild predator (IGP) on aphid traits. After the introduction of the aphids (N116 or Q1) followed by the parasitoid daughter into the microcosm, as explained above, an aphid lion second-instar larva was transferred into the microcosm, on a fine paintbrush, onto the soil a few minutes after the wasp was added. The daughters (parasitoid genotype) that arose from each of the sire × dam mating groupings were numbered and then split randomly into one of two groups: without IGP (*i.e*. IGP absent) or with IGP (*i.e*. IGP present). Once the microcosm set up was completed, they were sealed and placed back into the growth chamber for eleven days at 22-24°C with the 16h:8h photoperiod as above. The microcosms were randomised in the chamber and checked to ensure that they had enough water every other day. On the eleventh day, the microcosms were once again removed from the growth chamber, opened and the data were recorded. We recorded the total number of healthy aphids (non-mummified), the total number of mummies, and the distribution of the mummies within the microcosm (on versus off plant) Electronic supplementary material, figure S1. We were unable to create a fully factorial design with two aphid lineages and the presence or absence of a predator for each dam/sire combination. The differential survival in this multispecies system combined with the nature of the quantitative genetic design, and keeping all the parthenogenetic aphids at the same age led to unbalanced sample sizes for a given aphid lineage, which, nevertheless, is sufficiently powered for the number of replicates. Overall, there were 119 parasitoid daughters. Each group of daughters was split into two populations, with one (n = 73) being provided with pea aphid N116 as provision, while the other (n = 45) was provided with pea aphid Q1. Each of these two populations where further split into two groups, with one group exposed to intraguild predation by the aphid lion larva (n = 43, in the case of N116, and n = 15 in the case of Q1) and the other group not (n = 30, in the case of N116, and n = 30 in the case of Q1).

### Molecular Analysis

The healthy aphids in each microcosm were preserved in a cryogenic tube at −195°C, at The University of Manchester liquid nitrogen sample storage facility, for later molecular analysis. The identification of the bacterial symbionts in the two lineages of pea aphid consisted of two parts: 1) the use of diagnostic PCR to confirm the presence or absence of the defensive symbiont *H. defensa* and 2) 16s rRNA gene sequencing for the identification of other symbionts. The aphid samples were surface-sterilised [45], then the DNA was extracted using ‘Qiagen DNAEasy Blood and Tissue Kit’ small insect supplementary protocol [45]. As the aphids are soft-bodied insects, we altered step 1 of the protocol slightly, rather than freezing them in liquid nitrogen and grinding them up in a pestle and mortar they were homogenised in a sterile microcentrifuge tube using a sterile disposable microcentrifuge tube homogenisation pestle. In step 3, the lysis time was increased from three to six hours and the rest of the protocol was followed with no further modifications. Subsequently, we ran a Diagnostic PCR [46]; the PCR reactions were visualised on a 1% agarose gel with SafeView Nucleic Acid Stain with Bioline HyperLadder™ 1kb. Afterwards, we ran 16s Gene Sequencing for a total of 70 samples (35 Q1 and 35 N116), which were sent for sequencing using GATC Biotech’s T7 sequencing primers. Once we had received the sequence data both the vector sequences and the parts of the sequences that contained bases that were below the confidence threshold were removed. The sequences were then analysed using the NCBI ‘standard nucleotide BLAST’ (megablast) and the Nucleotide collection (nr/nt). The most closely related bacteria were selected based on the blast output and where they fall on the resulting distance tree of the results (Electronic supplementary materials, Molecular Analysis).

### Statistics

The data on the parasitoid genotype with and without IGP were pooled because this enabled us to investigate the influence of the IGP on the outcome of the parasitoid genotype effect on aphid fitness (in terms of immunity to the parasitoid) and the behaviour of the aphid lineages. All statistical analyses were conducted using R [47] via RStudio [48]. Firstly, we tested the effects of parasitoid and aphid genetic variability in the absence or presence of IGP on aphid immunity ratio (IR: the proportion of aphids that was non-mummified [unparasitoidised] after 11 days of exposure to the parasitoid genotype relative to the entire population of aphids [healthy and mummified] per aphid lineage per microcosm). A generalised linear mixed effect model (Model 1) was applied with Poisson family, R packages ‘car’ [49] and Ime4 [50]. The explanatory variables were the following fixed effects: (1) IGP (No, Yes), (2) aphid lineage (N116, Q1), (3) parasitoid intraspecific genetic variation effect (daughters’ identity as per their sire x dam grouping that was the product of the quantitative genetic design), (4) the interaction (parasitoid genotype x aphid lineage), (5) the interaction (IGP x parasitoid genotype), and (6) the interaction (IGP x aphid lineage). The microcosm was modelled as a random effect. Secondly, we analysed aphid behaviour as the proportion of aphid mummies off plant relative to the total number of mummies in the microcosm, using the explanatory variables (1-4) as in Model 1, in a generalised linear model (Model 2) with a quasiPoisson family due to non-normality of the count data, R package ‘multcomp’ [51].

## 3. Results and Discussion

In this study, we investigated the effects of genetic variation in a parasitoid provided with two aphid host conspecifics (N116 and Q1) having different life histories and biotypes, on host fitness and behaviour under intraguild predation. As a measure of fitness, we focussed on aphid immunity ratio (IR) that is the proportion of healthy aphids to the total population of healthy and parasitoidised individuals [mummified]), and host avoidance behaviour.

### Differences in immunity between the two aphid lineages (N116 and Q1)

In a pilot study, we had established that the two different clonal lineages are very different in the susceptibility to the parasitoid wasp. Based on the known effects of defensive endosymbionts, we hypothesised that the two lineages differed in the defensive endosymbiont community they host. We, therefore, conducted an assay of the endosymbionts, which revealed that, unlike the Q1 genotype, N116 harboured different endosymbionts known to confer immunity to parasitoidism by the wasp *A. ervi* (Electronic supplementary materials, Molecular Analysis). Of the 35 samples that were sequenced in the N116 clone, 26 were successful and contained a long enough sequence (590bp to 1112bp) to conduct a BLAST analysis. Of the 26 BLAST analysed samples, two were found to contain chimeric sequences and have been excluded, 13 samples matched with the known defensive secondary symbiont *H. defensa* (99.19% to 99.87% identity); a defensive secondary symbiont found throughout pea aphid lineages [52] and reported to provide immunity to parasitoidism by stopping the development of the *A. ervi* larva, and hence rescuing the aphid host [11,52]. The level of conferred immunity can vary substantially by different strains of *H. defensa* and the spread of the endosymbiont may rapidly increase, in experimental populations, with exposure to parasitoid wasps [53]. Variation in protection is further influenced by the presence or absence of infection of the bacteria with different bacteriophages called APSEs [53]. These bacteriophages are thought to encode putative toxins that function in the specific defence against *A. ervi* [14,52], which, however, we did not investigate in this study.

Furthermore, we also found nine samples were most closely related to *Fukatsuia symbiotica* (99% to 100% identity), previously referred to as the X-type or PAXS symbiont, that, when found in association with *H. defensa*, provides high levels of resistance to *A. ervi* [55,56]; *F. symbiotica* and *H. defensa* were previously reported in the N116 lineage [42]. Interestingly we also found that one sequence was most closely related to *Serratia symbiotica* (99% identity), another known symbiont of aphids that provides resistance against parasitoids [11,15,57,58]. *Serratia symbiotica* has not been reported in this lineage before. Given the lack of evidence of strong immunity in pea aphids by means of encapsulating parasitoid eggs [15,16], the immunity of N116 was dependent on the presence of this set of defensive endosymbionts [13]. Still, the presence of three defensive symbionts in the N116 lineage is unusual and further work is required to understand the significance of this finding. For Q1, of the 35 samples sent for sequencing 23 were received and of sufficient quality for BLAST analysis (420bp to 967bp). Here, 20 samples positively matched with the secondary symbiont *S. symbiotica* (99% to 100% identity) but no *H. defensa* was identified, while three samples identified the primary endosymbiont *Buchnera aphidicola* (Electronic supplementary materials, Molecular Analysis).

### The effects of intraspecific genetic variation in the parasitoid and the aphid on aphid immunity when intraguild predation occurs

Having established differences in endosymbiont community, we then proceeded to our full experiment in which we focussed on aphid immunity as defined above. As shown in figure 1, the overall average immunity ratio (IR) of N116 was ~65% in the absence of IGP that increased to 86% when IGP was present. By contrast, the average IR of Q1 was ~20% (IGP absent) that slightly increased to ~27 when the IGP was present. Thus, IR in N116 was 3.25 times higher than in the Q1 lineage without IGP, and ~3.2 times higher with IGP. The IR was significantly affected by aphid lineage (F_(1,58)_ = 28.5, P < 0.0001) and parasitoid genotype (F_(37,58)_ = 95.76, P < 0.0001), the interaction between aphid lineage and parasitoid genotype (F_(3,58)_ = 12.1, P = 0.007), and the interaction between IGP and parasitoid genotype (F_(15,58)_ = 37.23, P = 0.001); the IGP effect on its own was not significant (see also electronic supplementary materials, Table S1 for the model summary). We recorded total immunity (IR = 100%) to parasitoid genotype in 10 out of 30 cases for N116 versus only one case out of 30 for Q1 when IGP was absent, and 24 cases out of 43 for N116 versus only one case out of 15 for Q1 when IGP was present. Conversely, for lack of immunity (IR = 0%), there were six cases out of 30 for N116 versus only 16 cases out of 30 for Q1 when IGP was absent, and four cases out of 43 for N116 versus only nine cases out of 15 for Q1 when IGP was present. See electronic supplementary materials, figures S2 and S3 for further details.

**Figure 1.**
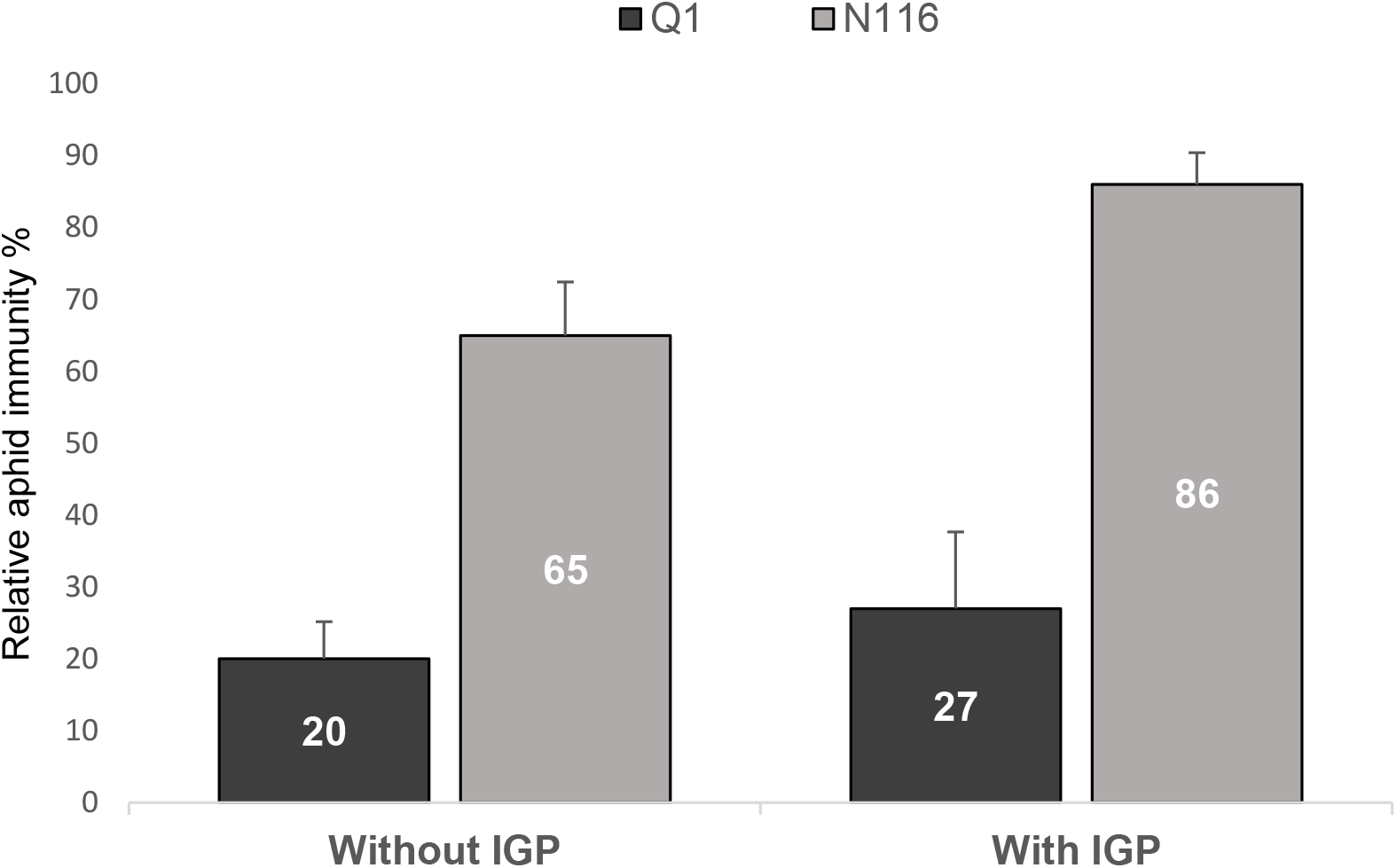
Aphid immunity to parasitoid subject to IGP. The means of aphid IR are proportionally shown per aphid lineage with and without IGP. Percentages of mummies recorded off the plant are shown for N116 (shown in grey) and Q1 (shown in black) pea aphids under exposure to the effect of the parasitoid genotype (daughters) in the absence of IGP (n = 60 parasitoid daughters [30 in the case of N116, and 30 in the case of Q1]) and the presence of IGP (n = 58 parasitoid daughters [43 in the case of N116, and 15 in the case of Q1]).

The parasitoid was less successful in sequestering aphids as puparia when the intraguild predator was present, and that was clearly pronounced in the N116 aphid lineage, which harboured defensive symbionts. The differences in the outcome of parasitoidism were influenced by the presence of the combination of defensive symbionts in N116 rather than in the Q1 lineage. This means, therefore, that the effect of the aphid ‘genotype’ in this work is more than the effect of the genotype alone, as it also includes the indirect factor of defensive endosymbiosis in association with the lineage. Isolates of both *S. symbiotica* and *H. defensa* have been shown to confer resistance to parasitoid wasps in the pea aphid, reducing successful parasitism by 23% and 42% accordingly [11,15]. Moreover, the occurrence of superinfected aphid clones (carrying multiple inherited symbionts), has been noted despite the apparent costs to aphid fecundity [57]. Aphids superinfected with *H. defensa* and *F. symbiotica* are known to have very high levels of resistance against *A. ervi*, up to 100% in some clones [42], and this explains the high levels of resistance in the N116 lineage. As such, the symbiosis, in this context, alters the outcome of the interaction between the parasitoid and the aphid host and thus should be considered as an important indirect ecological effect in this system [59]. We advocate that the indirect ecological effect influenced the outcome of the interspecific indirect genetic effect of the parasitoid on the reproductive success of its aphid hosts.

The strong and intimate interaction between the aphid host and its parasitoid may be influenced by genetic variation in the traits related to the interaction of the species involved, meeting one of the fundamental criteria for co-evolution in a host-parasitoid system [12]. At any rate, although the N116 pea aphid is one of the lineages with a known association with *H. defensa* [40,42], to the best of our knowledge, we are the first to empirically test the immunity in this lineage when an intraguild predator is present.

### Aphid altruistic mummification behaviour

Aphid mummification off plant, away from the healthy clonal population, has been interpreted as altruistic behaviour because it leads to an increased predation risk for parasitised aphids but a reduction in successfully eclosing parasitoid wasps [36,60]. Figure 2 shows that the average off-plant proportions of mummified aphids were almost identical in the case of N116 (~76%) with and without IGP. By contrast, for Q1 the average percentage was ~ 49% in the absence of IGP, and ~ 45% in presence of IGP. Parasitised N116 individuals mummified ~1.55 times more than Q1 when the IGP was absent, and ~1.69 times more in the presence of IGP. The proportion of mummies off plant were significantly affected by aphid lineage (LR_χ^2^(1,49)_ = 7.051, P = 0.008) and, marginally, parasitoid genotype (F_(28,49)_ = 40.62, P = 0.058), but the effect of their interaction was not significant, nor was the effect of IGP. See electronic supplementary materials, Table S2 for the model summary. These results suggest that N116 (relatively highly immune to parasitoid attack owing to the defensive endosymbiont) showed a consistent propensity to desert the host plant when parasitised. Yet, the ecological effect of IGP on such a propensity was negligible. Comparatively, Q1 (with inferior immunity due to differences in their defensive symbiont communities, see above) showed less altruistic behaviour when the aphid lion was present.

**Figure 2.**
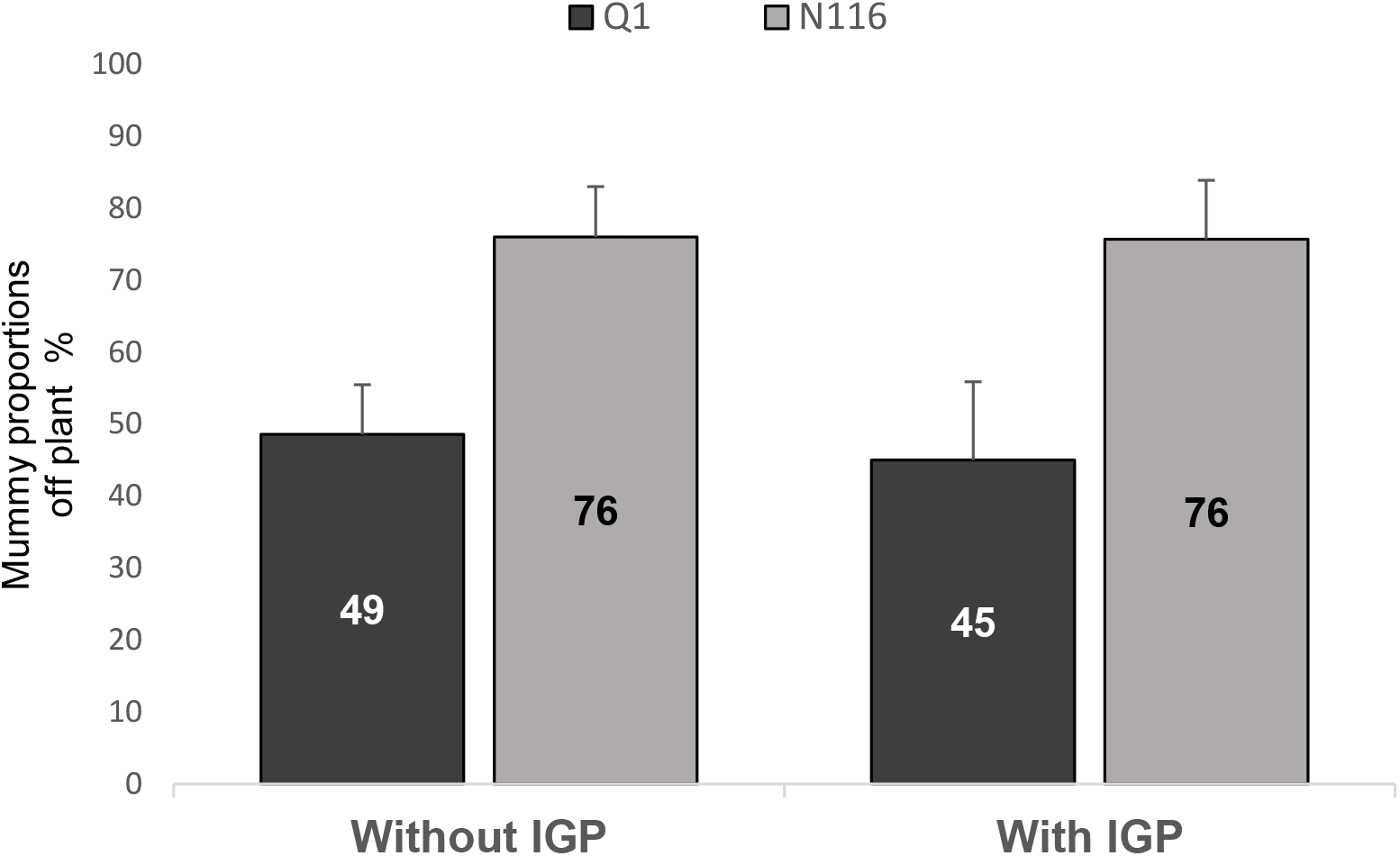
Proportions of mummies off the plant. Percentages of mummies (means) recorded off the plant are shown for N116 and Q1 pea aphids under exposure to the effect of the parasitoid genotype (daughters) in the absence of IGP (n = 49 parasitoid daughters [20 in the case of N116, and 29 in the case of Q1]) and the presence of IGP (n = 33 parasitoid daughters [19 in the case of N116, and 14 in the case of Q1]).

Under the life-dinner principle [61], changes in aphid population and altruistic behaviour may lead to changes in the parasitoid host-manipulative tactics and virulence, such that a decreased aphid altruistic behaviour may reduce the parasitoid loss inflicted by an intraguild predator (which shares the aphid as prey with the parasitoid). Thus, parasitoid wasps alter the behaviour [21] as well as the internal environment of the parasitised aphid to make it more favourable for wasp development and survival [62]. Our findings show that lineage-specific factors, including the absence or presence of defensive endosymbiosis influence on the location of the mummies. This indicates that the response of specific lineages of the pea aphid to the intraspecific genetic variation effects and interspecific indirect genetic effects of its parasitoid is also dependent on the aphid within-species genetic variability as well. Having more aphid mummies (wasp puparia) farther from the core of the mother clone is assumed to increase aphid inclusive fitness, but this altruistic change in mummy position is likely to be cost-sensitive and context-dependent [21,60,63–65]. It is worthy of note here that parasitoid wasps may be able to differentiate between infected and uninfected aphids, thought to be the result of a decreased production of a major component of the aphid alarm pheromone, *trans*-β-farnesene (EBF) [62]. The alarm pheromone is secreted from cornicles when the aphids are attacked, and when aphids detect this pheromone they move away from the source, with some even dropping from the plant altogether [62]. This potential of *A. ervi* to differentiate between aphids infected with *H. defensa* and those that are not is demonstrated by an increased occurrence of superparasitism in the infected aphids. Superparasitism occurs when more than one egg is oviposited into the same aphid host and, under normal conditions, this behaviour is usually considered to be maladaptive as it results in siblicide [66]. Interestingly, the presence of *H. defensa* in a host aphid may have further implications for the plant-aphid-parasitoid system as it alters the behaviour of the parasitoids [62,14]. Vorburger and Rouchet (2016) [67] suggested that there may be selection for local adaption by parasitoids to certain strains of *H. defensa*, but this remains in need of further investigation [67]. This implies that the interaction between the aphid (including the defensive symbiosis) and the parasitoid is highly context-dependent as shown in our study. Moreover, *H. defensa* is also implicated in changing aphid defensive behaviour against parasitoids [68] and in attenuating the release of herbivore-induced plant volatiles that attract parasitoid wasps [69]. This further highlights the importance of symbionts in the interactions between species [14,69] such that defensive symbionts are reported to have far-reaching ecological effects on aphid-parasitoid communities [70]. The rate of evolution of host resistance to parasitoids, as well as the infectivity (virulence) of parasitoids will be subject to the impacts of internal defensive symbionts [65,72,71] and external factors (e.g. intraguild predators) [32,33]. Altogether, these are constituents of ongoing evolutionary arms-race [31,37–39] that will depend on the levels of variation present in the populations and the associated fitness costs of the involved traits [65,72]. This is in line with the extensive effects of intra-specific genetic variation of one species on other species beyond the individual or population levels [18,19].

Our study has demonstrated the complex nature of the interaction between two lineages of a scientifically as well as economically important agricultural pest and the genotype of its parasitoid subject to the effects of intraguild predation by aphid lion. Our findings imply that having defensive endosymbiosis may contribute to aphid survival and reactions to differential parasitoid virulence that appear to be context-dependent. The influence of the presence of the intraguild predator varied across parasitoid genotypes and aphid lineages. We demonstrate the need to consider the effects of intra-specific genetic variation in host-parasitoid systems together with the ecological effects brought about by defensive endosymbiosis and other natural enemies of the aphid across trophic levels. This will help untangle the complexity of these interactions and hence design effective biological controls in agro-ecosystems.

## Data accessibility

The data analysed in this study are provided in the Figshare repository: https://figshare.com/s/96df54283ac09ebd39cd

## Gene Bank accession numbers

N116 aphid endosymbionts: MW979375 to MW979398

Q1 aphid endosymbionts: MW971996 to MW972018

## Acknowledgements

We would like to thank Marco Giorda for help with data collection, and Dr Jon Pittman for general advice, and Reuben Margerison for his assistance with the 16s sequence analysis.

