## Supplementary material for "Symbiont-conferred immunity interacts with the effects of parasitoid genotype and intraguild predation to shape pea aphid immunity in a clone-specific fashion"

**Electronic Supplementary Materials**

19

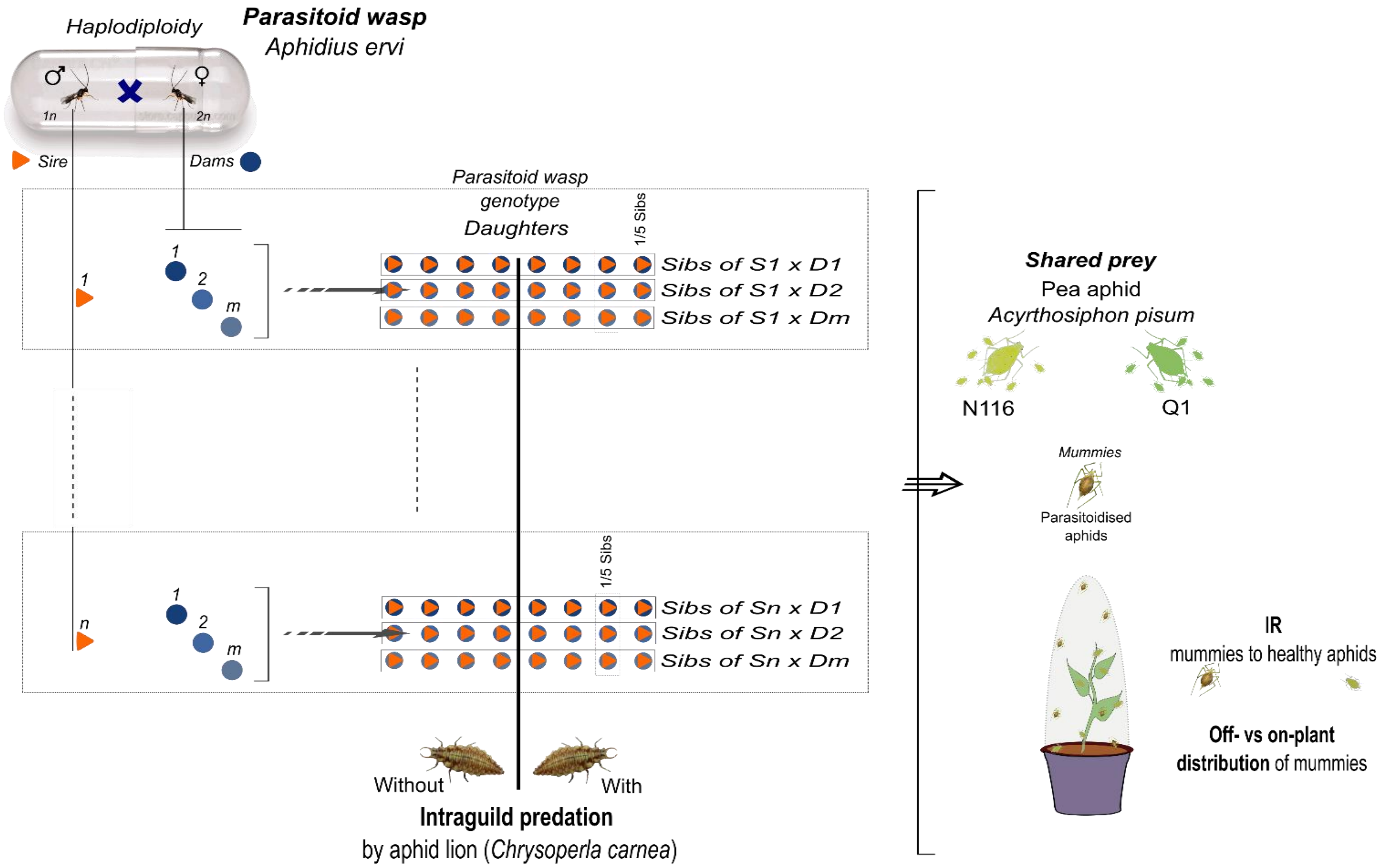
Haplodiploidy
Parasitoid wasp
Aphidius ervi
♂
♀
1n
2n
Sire
Dams
Parasitoid wasp
genotype
Daughters
1/5 Sibs
Sibs of S1 x D1
Sibs of S1 x D2
Sibs of S1 x Dm
Sibs of Sn x D1
Sibs of Sn x D2
Sibs of Sn x Dm
Shared prey
Pea aphid
Acyrthosiphon pisum
N116
Q1
Mummies
Parasitoidised aphids
IR
mummies to healthy aphids
Off- vs on-plant distribution of mummies
Without
With
Intraguild predation
by aphid lion (Chrysoperla carnea)

**Figure S1. Experimental design.** The diagram shows the full experimental design, with n sires being mated to at least three dams (D1 – D*m*). The sire × dam mating groupings produced the intraspecific genetic variability in the parasitoid (genotype/daughters [sibs, and half-sibs]). Overall, there were 119 parasitoid daughters. Each group of daughters of the (S1 – S*n*) combinations was then split into two populations, with one (n = 73) being provided with pea aphids of the N116 lineage as a provision, while the other (n = 45) provided with pea aphids of the Q1 lineage. Each of these populations where further split into two groups, with one group exposed to intraguild predation by the aphid lion larva (n = 43, in the case of N116, and n = 15 in the case of Q1) and the other group not (n = 30, in the case of N116, and n = 30 in the case of Q1). The effects of parasitoid and aphid genetic variability (with and without the aphid lion) on aphid immunity ratio (IR) were investigated in microcosms. IR is the proportion of healthy aphids (non-mummified *i.e.* unparasitoidised) after 11 days of exposure to the parasitoid genotype relative to the entire population of aphids (healthy and mummified) per aphid lineage.

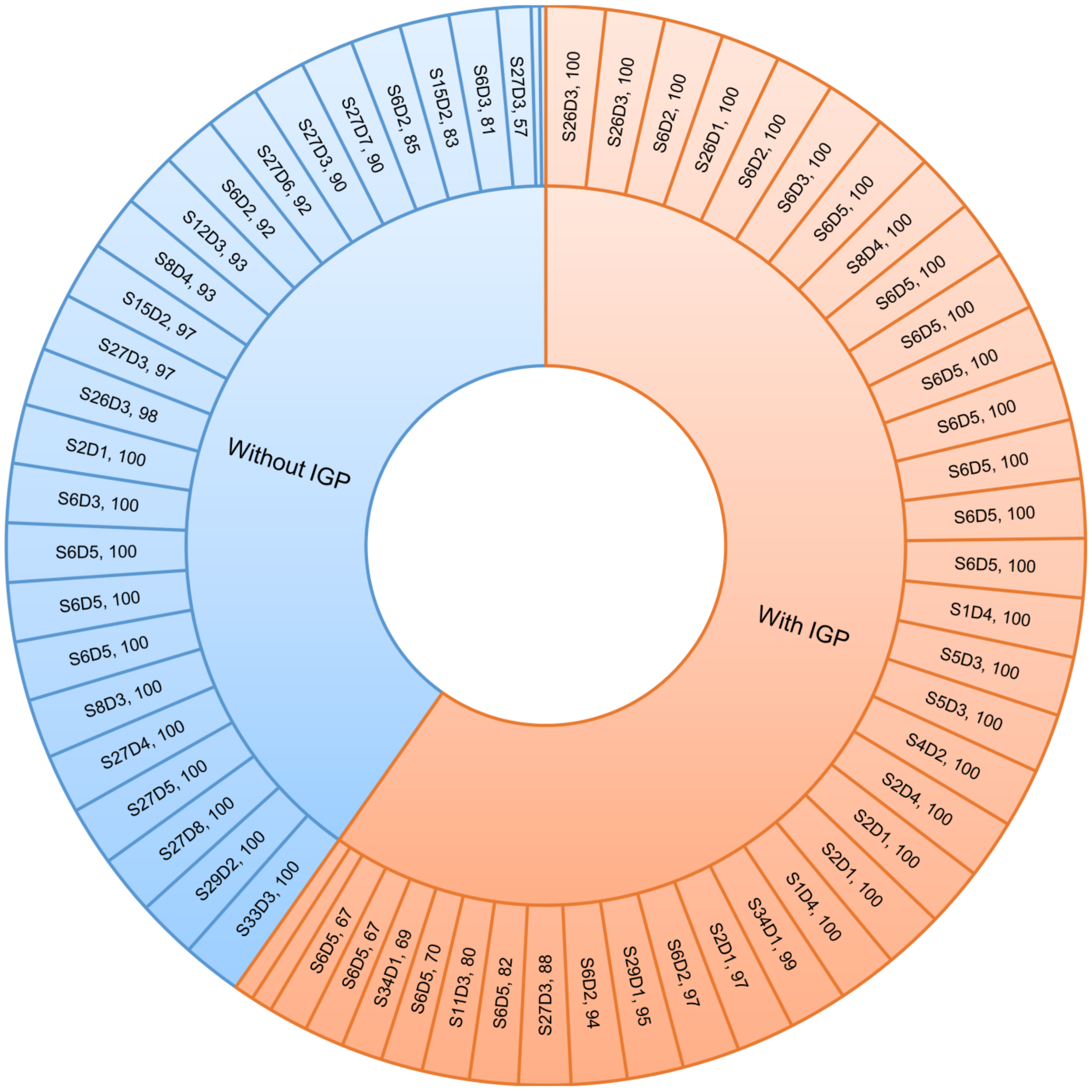
Without IGP
With IGP
S27D3, 57
S6D3, 81
S15D2, 83
S6D2, 85
S27D7, 90
S27D3, 90
S27D6, 92
S6D2, 92
S12D3, 93
S8D4, 93
S15D2, 97
S27D3, 97
S26D3, 98
S2D1, 100
S6D3, 100
S6D5, 100
S6D5, 100
S6D5, 100
S8D3, 100
S27D4, 100
S27D5, 100
S27D8, 100
S29D2, 100
S33D3, 100
S26D3, 100
S26D3, 100
S6D2, 100
S26D1, 100
S6D2, 100
S6D3, 100
S6D5, 100
S8D4, 100
S6D5, 100
S6D5, 100
S6D5, 100
S6D5, 100
S6D5, 100
S6D5, 100
S6D5, 100
S1D4, 100
S5D3, 100
S5D3, 100
S4D2, 100
S2D4, 100
S2D1, 100
S2D1, 100
S1D4, 100
S34D1, 99
S2D1, 97
S6D2, 97
S29D1, 95
S6D2, 94
S27D3, 88
S6D5, 82
S11D3, 80
S6D5, 70
S34D1, 69
S6D5, 67
S6D5, 67

**Figure S2. Detailed N116 immunity to parasitoid genotype with and without IGP.** This ‘sunburst’ infographic illustrates the detailed immunity ratio (IR) in the N116 pea aphid lineage to the parasitoid daughters (genotype effect). IR in the microcosm is the proportion of healthy aphids (non-mummified *i.e.* unparasitoidised) after 11 days of exposure to the parasitoid genotype relative to the entire population of aphids (healthy and mummified) per aphid lineage. The inner circle represents the absence of IGP (Without IGP) or its presence (With IGP), while the outer circle represents the parasitoid daughters including the corresponding IR value (%) separated by a comma. There were 73 parasitoid daughters in total as the result of the quantitative genetic design (with IGP, n = 30 parasitoid daughters, and without IGP, n = 43 parasitoid daughters).

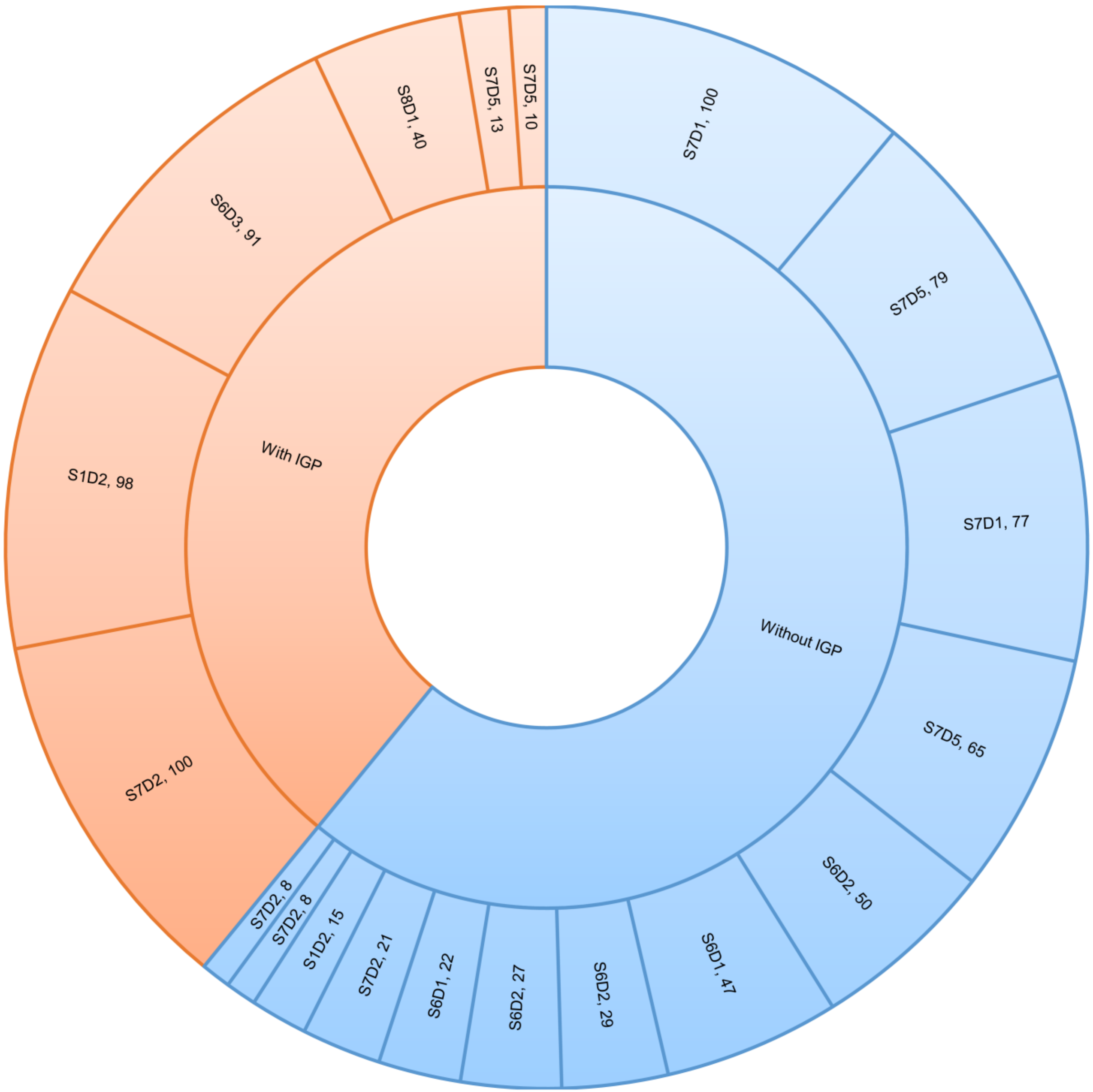
With IGP
Without IGP
S7D5, 10
S7D5, 13
S8D1, 40
S6D3, 91
S1D2, 98
S7D2, 100
S7D1, 100
S7D5, 79
S7D1, 77
S7D5, 65
S6D2, 50
S6D1, 47
S6D2, 29
S6D2, 27
S6D1, 22
S7D2, 21
S1D2, 15
S7D2, 8
S7D2, 8

40

**Figure S3. Detailed Q1 immunity to parasitoid genotype with and without IGP.** This ‘sunburst’ infographic illustrates the detailed immunity ratio (IR) in the Q1 pea aphid lineage to the parasitoid daughters (genotype effect). IR in the microcosm is the proportion of healthy aphids (non-mummified *i.e.* unparasitoidised) after 11 days of exposure to the parasitoid genotype relative to the entire population of aphids (healthy and mummified) per aphid lineage. The inner circle represents the absence of IGP (Without IGP) or its presence (With IGP), while the outer circle represents the parasitoid daughters including the corresponding IR value (%) separated by a comma. There were 45 parasitoid daughters in total as the result of the quantitative genetic design (with IGP, n = 30 parasitoid daughters 30, and without IGP, n = 15 parasitoid daughters).

### Molecular Analysis

**Table 1. Primers used in this study.** The table shows the target symbiont, the respective target genes with their primer names, followed by the sequence (5’-3’) and the references given below the table.

| Target symbiont | Target gene | Primer name | Sequence (5’-3’) | References* |
|---|---|---|---|---|
| **Primers used in diagnostic 16s PCR** | | | | |
| *H. defensa* | 16s rRNA | 10F | AGTTTGATCATGGCTCAGATTG | 1 |
| *H. defensa* | 16s rRNA | 16s- T419R | AAATGGTATTSGCATTTATCG | 1 |
| **Primers used in PCR reactions for 16s gene sequencing** | | | | |
| Universal | 16s rRNA | fD1 | AGAGTTTGATCCTGGCTCAG | 2 |
| Universal | 16s rRNA | rP2 | ACGGCTACCTTGTTACGACTT | 2 |

* References: 1) Henry, L., Peccoud, J., Simon, J., Hadfield, J., Maiden, M., Ferrari, J. and , H., 2013. Horizontally transmitted symbionts and host colonization of ecological niches. Current Biology, 23(17), pp.1713-1717. 2) Weisburg, W., Barns, S., Pelletier, D. and Lane, D., 1991. 16S ribosomal DNA amplification for phylogenetic study. Journal of Bacteriology, 173(2), pp.697-703.

**Gel extraction**

Once the universal 16s PCR reactions were completed [*95°C 5 mins, (95°C 30s, 60°C 30s, 72°C 30 sec) x 40 and a final extension of 72°C 7 mins*] they were visualised on a 1% agarose gel with SafeView Nucleic Acid Stain with Bioline HyperLadder™ 1kb. The gel bands were cut from the gel under UV light using a sterile scalpel and extracted using the Qiagen QIAquick gel extraction kit.

**Ligation and transformation**

The ligation of the samples into a plasmid was conducted using the Promega ‘pGEM®-T Easy Vector System 1’ as per the manufacturer's protocol and set up in the following reaction and left overnight at 4°C.

The transformation reaction used XL1-Blue competent cells (Agilent Technologies) and the manufactures ‘transformation protocol’ with some slight alterations. We added 2 µl of the ligated plasmid sample to the aliquot of cells (Step 5). We also used LB media instead of the suggested SOC media (step 9) and the LB-ampicillin plates were made by adding 15g of agar and 25g of LB medium to 1L of MiliQ water and mixing well. The LB agar was then autoclaved and left to cool to below 55°C. When the mixture reached the correct temperature, it was taken to a laminar flow cabinet and the filter-sterilized ampicillin was added (1 ml of a 100mg/ml solution). The agar is then mixed again and poured into Petri dishes and left to set in the laminar flow, under flame. For colour screening, 100µl of 100 mM IPTG and 20µl of 50 mg/ml X-gal was pipetted onto the agar and spread evenly (step 10). The plates were left for 48 hours for the colonies to develop as they were too small after the suggested 17 hours. After 48 hours of incubation at 37°C, the colonies were incubated at 4°C for 2 hours to enhance the colours. The growth of any bacteria that did not take up the plasmid would be inhibited by the antibiotic and if a colony formed and contained the plasmid but not the 16s gene we had tried to insert, then the colonies grow on the plates and appear blue in colour. Moreover, if the colonies contain the plasmid with the 16s gene insert, they appear white, making the colonies easy to sample. The colonies were sampled by touching them with a pipette tip and the tip was dropped into a falcon tube that contained 3ml of sterile LB-ampicillin media (same concentration of antibiotic as the plates). The sampled colonies were incubated at 37°C, with shaking at 225-250 rpm, overnight. The next day the plasmids were extracted from the sampled colonies using the Qiagen, QIAprep® spin miniprep kit and quick-start protocol and eluted with nuclease-free water.

**Sample preparation for sequencing**

Before the plasmids were sent for sequencing, 15 samples of each aphid line were digested with the restriction enzyme EcoR1. This enzyme removes the insert sequence from the plasmid we had used and enabled us to visually confirm its presence before sequencing. After digestion, when the samples were run on a 1% agarose gel, they showed two clear bands, one for the plasmid and a band of a similar size to the insert.

To prepare the samples for sequencing their concentration was checked on a Thermo Scientific™ nanodrop™ 2000 spectrophotometer. GATC Biotech AG, London UK, (now Eurofins GATC) recommended a DNA concentration of between 30 – 80 ng/µl for their plasmid 'Supreme run' Sanger sequencing. Those samples that had a concentration higher than recommended were then diluted, with nuclease-free water, to a final concentration of 60ng/µl. A total of 70 samples (35 Q1 and 35 N116) were sequenced using GATC Biotech's T7 sequencing primers.

**Analysis of sequencing data**

Sequences have been edited to remove pGEM-T Easy Vector sequences and any parts of the sequence that contained bases that did not pass the confidence threshold.

**N116 aphid symbiont BLAST analysis**

Trimmed 'N116' clone *Acyrthosiphon pisum* symbiont 16s rRNA gene sequences used in the blast analysis shown in FASTA format. GenBank Accession numbers: MW979375 to MW979398

**> Seq1 [organism=Candidatus *Hamiltonella defensa*] 16S ribosomal RNA gene [host=*Acyrthosiphon pisum* isolate N116]**
AGAGTTTGATCCTGGCTCAGATTGAACACTGGTGGCAGGCCTAACACATGCAAGTCGAGCGGCATCGAGTGAGCGCAGTTTACTGAGTTCATGTCGGCG
AGCGGCGGACGGGTGAGTAAAGTCTGGGAATCTGGCCGAAGGAGGGGGATAACTGCTGGAAACGGCAGCTAATACCGCATGAAGTCGCGAGACCAAAG
TGGGGGACCTTCGGGCCTCACGCCTTCGGATGAGCCCAGATGAGATTAGCTGGTAGGTAAGGTAAGGGCTTACCTAGGCGACGATCTCTAGCGGGTCT
GAGAGGATAGCCCGCCACACTGGAACTGAGACACGGTCCAGACTCCTACGGGAGGCAGCAGTGGGGAATATTGCACAATGGGCGAAAGCCTGATGCAG
CCATGCCACGTGTGTGAAGAAGGCCTTCGGGTTGTAAAGCACTTTCAGCGAGGAGGAAGCGATAAATGCGAATACCATTTATTTTTGACGTTACTCGCAG
AAGAAGCACCGGCTAACTCCGTGCCAGCAGCCGCGGTAATACGGAGGGTGCGAGCGTTGATCGGAATAACTGGGCGTAAAGGGCATGTAGGCGGTGAG
TTAAGTCAGATGTGAAATCCCCGAGCTCAACTTGGGAATGGCATTTGAAACTGGGTCGCTAGAGTTTTCTAGAGGGGGGTAGAATTCCAGGTGTAGCGGT

GAAATGCGTAGATATCTGGAGGAATACCGGTGGCGAAGGCGGCCCCCTGGAGAAAGACTGACGCTGAGGTGCGAAAGCGTGGGGAGCAAACAGGATTA
GATACCCTGGTAGTCCACGCTGTAAACGATGTCGATTTGGAGGTTGCGGTCTTGAACTGTGGCGTCCGGAGCTAACGCG

**> Seq2 [organism=Candidatus *Fukatsuia symbiotica*] 16S ribosomal RNA gene [host=*Acyrthosiphon pisum* isolate N116]**
AGAGTTTGATCCTGGCTCAGATTGAACGCTGGCGGCAGGCCTAACACATGCAAGTCGAGCGGCATCGGGAAGGTAGCTTGCTATCTTTGCCGGCGAGC
GGCGGACGGGTGAGTAAAGTCTGGGGATCTGCCTGATGGAGGGGGATAACTACTGGAAACGGTAGCTAATACCGCATGATGTTACGCGACCAAAGCGG
GGGACCTCCGGGCCTCGCGCCATCAGATGAACCCAGATGGGATTAGCTAGTAGGAGAGGTAATGGCTCCCCTAGGCGACGATCCCTAGCTGGTCTGAG
AGGATAACCAGCCACACTGGAACTGAGAGACGGTCCAGACTCCTACGGGAGGCAGCAGTGGGGAATATTGCACAATGGGCGCAAGCCTGATGCAGCCA
TGCCGCGTGTGTGAAGAAGGCCTTCGGGTTGTAAAGCACTTTCAGCGAGGAGGAATGAAGCAATGCAAAGAGTGTTGCTAATGGACGTTACTCGCAGAA
GAAGCACCGGCTAACTCCGTGCCAGCAGCCGCGGTAATACGGAGGGTGCGAGCGTTAATCGGAATTACTGGGCATAAAGGGCACGTAGGCGGTTTCTT
AAGTCAGATGTGAAATCCCCGAGCTTCACTTGGGAACGGCATTTGAAACTGAGAGTCTAGAGTTTTGTAGAGGGGGGTAGAATTCCAGGTGTAGCGGTGA
AATGCGTAGATATCTGGAGGAATACCGGTGGCGAAGGCGGCCCCCTGGACAGAGACTGACGCTGAGGTGCGAAAGCGTGGGTAGCAAACAG

**> Seq3 [organism=Candidatus *Fukatsuia symbiotica*] 16S ribosomal RNA gene [host=*Acyrthosiphon pisum* isolate N116]**
AGAGTTTGATCCTGGCTCAGATTGAACGCTGGCGGCAGGCCTAACACATGCAAGTCGAGCGGCATCGGGAAGGTAGCTTGCTATCTTTGCCGGCGAGC
GGCGGACGGGTGAGTAAAGTCTGGGGATCTGTCTGATGGAGGGGGATAACTACTGGAAACGGTAGCTAATACCGCATGATGTTACGCGACCAAAGCGG
GGGACCTCCGGGCCTCGCGCCATCAGATGAACCCAGATGGGATTAGCTAGTAGGAGAGGTAATGGCTCCCCTAGGCGACGATCCCTAGCTGGTCTGAG
AGGATAACCAGCCACACTGGAACTGAGAGACGGTCCAGACTCCTACGGGAGGCAGCAGTGGGGAATATTGCACAATGGGCGCAAGCCTGATGCAGCCA
TGCCGCGTGTGTGAAGAAGGCCTTCGGGTTGTAAAGCACTTTCAGCGAGGAGGAATGAAGCAATGCAAAGAGTGTTGCTAATGGACGTTACTCGCAGAA
GAAGCACCGGCTAACTCCGTGCCAGCAGCCGCGGTAATACGGAGGGTGCGAGCGTTAATCGGAATTACTGGGCGTAAAGAGCACGTAGGCGGTTTCTT
AAGTCAGATGTGAAATCCCCGAGCTTCACTTGGGAACGGCATTTGAAACTGAGAGTCTAGAGTTTTGTAGAGGGGGGTAGAATTCCAGGTGTAGCGGTGA
AATGCGTAGATATCTGGAGGAATACCGGTGGCGAAGGCGGCCCCCTGGACAGAGACTGACGCTGAGGTGCGAAAGCGTGGGTAGCAAACAGGATTAGA
TACCCTGGTAGTCCACGCTGTAAACGATGTCGATTTGTAGGTTGTGGTTATAAACTGTGGCTTGCGGAGCAAACGCGTTAAATCGACCGCCTGGGGAGTA
CGGCCGCAAGGTTAAAACTCAAATGAATTGACGGGGGCCCGCACAAGCGGTGGAGCATGTGGTTTAATTCGATGCCACGCGAAGAACCTTACCTACTCT
TGACATCCAGAGG

**> Seq4 [organism=Candidatus *Hamiltonella defensa*] 16S ribosomal RNA gene [host=*Acyrthosiphon pisum* isolate N116]**
AGAGTTTGATCCTGGCTCAGGTTGAACACTGGTGGCAGGCCTAACACATGCAAGTCGAGCGGCATCGAGTGAGCGCAGTTTACTGAGTTCATGTCGGCG
AGCGGCGGACGGGTGAGTAAAGTCTGGGAATCTGGCCGAAGGAGGGGGATAACTGCTGGAAACGGCAGCTAATACCGCATGAAGTCGCGAGACCAAAG
TGGGGGACCTTCGGGCCTCACGCCTTCGGATGAGCCCAGATGAGATTAGCTGGTAGGTAAGGTAAAGGCTTACCTAGGCGACGATCTCTAGCGGGTCT
GAGAGGATAGCCCGCCACACTGGAACTGAGACACGGTCCAGACTCCTACGGGAGGCAGCAGTGGGGAATATTGCACAATGGGCGAAAGCCTGATGCAG
CCATGCCACGTGTGTGAAGAAGGCCTTCGGGTTGTAAAGCACTTTCAGCGAGGAGGAAGCGATAAATGCGAATACCATTTATTTTTGACGTTACTCGCAG
AAGAAGCACCGGCTAACTCCGTGCCAGCAGCCGCGGTAATACGGAGGGTGCGAGCGTTAATCGGAATAACTGGGCGTAAAGGGCATGTAGGCGGTGAG
CTAAGTCAGATGTGAAATCCCCGAGCTCAACTTGGGAATGGCATTTGAAACTGGGTCGCTAGAGTTTTCTAGAGGGGGGTAGAATTCCAGGTGTAGCGGT
GAAATGCGTAGATATCTGGAGGAATACCGGTGGCGAAGGCGGCCCCCTGGAGAAAGACTGACGCTGAGGTGCGAAAGCGTGGGGAGCAAACAGGATTA
GATACCCTGGTAGTCCACGCTGTAANCGATGTCGATTTGGAGGTTGCGGTCTTGAACTGTGGCGTCCGGAGCTAACGCGTTAAATCGACCGCCTGGGGA
GTACGGCCGCAAGGTTAAAACTCAAATGAATTGACGGGGGCCCGCACAAGCGGTGGAGCATGTGGTTTAATTCGATGCAACGCGAA

**> Seq5 [organism=Candidatus *Fukatsuia symbiotica*] 16S ribosomal RNA gene [host=*Acyrthosiphon pisum* isolate N116]**
AGAGTTTGATCCTGGCTCAGATTGAACGCTGGCGGCAGGCCTAACACATGCAAGTCGAGCGGCATCGGGAAGGTAGCTTGCTATCTTTGCCGGCGAGC
GGCGGACGGGTGAGTAAAGTCTGGGGATCTGCCTGATGGAGGGGGATAACTACTGGAAACGGTAGCTAATACCGCATGATGTTACGCGACCAAAGCGG
GGGACCTCCGGGCCTCGCGCCATCAGATGAACCCAGATGGGATTAGCTAGTAGGAGAGGTAATGGCTCCCCTAGGCGACGATCCCTAGCTGGTCTGAG
AGGATAACCAGCCACACTGGAACTGAGAGACGGTCCAGACTCCTACGGGAGGCAGCAGTGGGGAATATTGCACAATGGGCGCAAGCCTGATGCAGCCA
TGCCGCGTGTGTGAAGAAGGCCTTCGGGTTGTAAAGCACTTTCAGCGAGGAGGAATGAAGCAATGCAAAGAGTGTTGCTAATGGACGTTACTCGCAGAA
GAAGCACCGGCTAACTCCGTGCCAGCAGCCGCGGTAATACGGAGGGTGCGAGCGTTAATCGGAATTACTGGGCGTAAAGGGCACGTAGGCGGCTTCTT
AAGTCAGATGTGAAATCCCCGAGCTTCACTTGGGAACGGCATTTGAAACTGAGAGTCTAGAGTTTTGTAGAGGGGGGTAGAATTCCAGGTGTAGCGGTGA
AATGCGTAGATATCTGGAGGAATACCGGTGGCGAAGGCGGCCCCCTGGACAGAGACTGACGCTGAGGTGCGAAAGCGTGGGTAGCAANCAGGATTAGA
TACCCTGGTAGTCCACGCTGTAAACGATGTCGATTTGTAGGTTGTGGTTATAAACTGTGGCTTGCG

**> Seq6 [organism= Candidatus *Hamiltonella defensa*] 16S ribosomal RNA gene [host=*Acyrthosiphon pisum* isolate N116]**
AGAGTTTGATCCTGGCTCAGATTGAACACTGGTGGCAGGCCTAACACATGCAAGTCGAGCGGCATCGAGTGAGCGCAGTTTACTGAGTTCATGTCGGCG
AGCGGCGGACGGGTGAGTAAAGTCTGGGAATCTGGCCGAAGGAGGGGGATAACTGCTGGAAACGGCAGCTAATACCGCATGAAGTCGCGAGACCAAAG
TGGGGGACCTTCGGGCCTCACGCCTTCGGATGAGCCCAGATGAGATTAGCTGGTAGGTAAGGTAAAGGCTTACCTAGGCGACGATCTCTAGCGGGTCT
GAGAGGATAGCCCGCCACACTGGAACTGAGACACGGTCCAGACTCCTACGGGAGGCAGCAGTGGGGAATATTGCACAATGGGCGAAAGCCTGATGCAG
CCATGCCACGTGTGTGAAGAAGGCCTTCGGGTTGTAAAGCACTTTCAGCGAGGAGGAAGCGATAAATGCGAATACCATTTATTTTTGACGTTACTCGCAG
AAGAAGCACTGGCTAACTCCGTGCCAGCAGCCGCGGTAATACGGAGGGTGCGAGCGTTAATCGGAATAACTGGGCGTAAAGGGCATGTAGGCGGTGAG
TTAAGTCAGATGTGAAATCCCCGAGCTCAACTTGGGAATGGCATTTGAAACTGGGTCGCTAGAGTTTTCTAGAGGGGGGTAGAATTCCAGGTGTAGCGGT
GAAATGCGTAGATATCTGGAGGAATACCGGTGGCGAAGGCGGCCCCCTGGAGAAAGACTGACGCTGAGGTGCGAAAGCGTGGGGAGCAAACAGGATTA
GATACCCTGGTAGTCCACGCTGTAAACGATGTCGATTTGGAGGTTGCGGTCTTGAACTGTGGCGTCCGGAGCTAACGCGTTAAATCGACCGCCTGGGGA
GTACGGCCGCAAGGTTAAAACTCAAATGAATTGACGGGGGCCCGCACAAGCGGTGGAGCATGTGGTTTAATTCGATGCAACGC

**>Seq7 [organism=Candidatus *Fukatsuia symbiotica*] 16S ribosomal RNA gene [host=*Acyrthosiphon pisum* isolate N116]**
GGTTACCTTGTTACGACTTCACCCCAGTCATGGTTCACAAAGTGGTAAGCGCCATCCCAAAGGTTAAGCTACCTACTTCTTTTGCAAAACATTCCCATGGT
GTGACGGGCGGTGTGTACAAGGCCCGGGAACGTATTCACCGTAGCATTCTGATCCACGATTACTAGCGATTCCGACTTCATGGAGTCGAGTTGCAGACT
CCAATCCGGACTACGACGTACTTTATGAGGTCCGCTCACCCTCGCAGGCTCGCTTCTCTTTGTATACGCCATTGTAGCACGTGTGTAGCCCTACTCGTAA
GGGCCATGATGACTTGACGTCATCCCCACCTTCCTCCGGTTTATCACCGGCAGTCTCTCTTGAGTTCCCACCTCTACGTGCTGGCAACAAAAGATAAGGG
TTGCGCTCGTTGCGGGACTTAACCCAACATTTCACAACACGAGCTGACGACAGCCATGCAGCACCTGTCTCAAAGCTCCCCGAAGGGCACGTCAACATC
TCTGTCGACTCCTCTGGATGTCAAGAGTAGGTAAGGTTCTTCGCGTTGCATCGAATTAAACCACATGCTCCACCGCTTGTGCGGGCCCCCG

**> Seq9 [organism=Candidatus *Hamiltonella defensa*] 16S ribosomal RNA gene [host=*Acyrthosiphon pisum* isolate N116]**
AGAGTTTGATCCTGGCTCAGATTGAACACTGGTGGCAGGCCTAACACATGCAAGTCGAGCGGCATCGAGTGAGCGCAGTTTACTGAGTTCATGTCGGCG
AGCGGCGGACGGGTGAGTAAAGTCTGGGAATCTGGCCGAAGGAGGGGGATAACTGCTGGAAACGGCAGCTAATACCGCATGAAGTCGCGAGACCAAAG
TGGGGGACCTTCGGGCCTCACGCCTTCGGATGAGCCCAGATGAGATTAGCTGGTAGGTAAGGTAAAGGCTTACCTAGGCGACGATCTCTAGCGGGTCC
GAGAGGATAGCCCGCCACACTGGAACTGAGACACGGTCCAGACTCCTACGGGAGGCAGCAGTGGGGAATATTGCACAATGGGCGAAAGCCTGATGCAG
CCATGCCACGTGTGTGAAGAAGGCCTTCGGGTTGTAAAGCACTTTCAGCGAGGAGGAAGCGATAAATGCGAATACCATTTATTTTTGACGTTACTCGCAG
AAGAAGCACCGGCTAACTCCGTGCCAGCAGCCGCGGTAATACGGAGGGTGCGAGCGTTAATCGGAATAACTGGGCGTAAAGGGCATGTAGGCGGTGAG
TTAAGTCAGATGTGAAATCCCCGAGCTCAACTTGGGAATGGCATTTGAAACTGGGTCGCTAGAGTTTTCTAGAGGGGGGTAGAATTCCAGGTGTAGCGGT
GAAATGCGTAGATATCTGGAGGAATACCGGTGGCGAAGGCGGCCCCCTGGAGAAAGACTGACGCTGAGGTGCGAAAGCGT

**> Seq10 [organism=Candidatus *Hamiltonella defensa*] 16S ribosomal RNA gene [host=*Acyrthosiphon pisum* isolate N116]**
AGAGTTTGATCCTGGCTCAGATTGAACACTGGTGGCAGGCCTAACACATGCAAGTCGAGCGGCATCGTGTGAGCGCAGTTTACTGAGTTCATGTCGGCG
AGCGGCGGACGGGTGAGTAAAGTCTGGGAATCTGGCCGAAGGAGGGGGATAACTGCTGGAAACGGCAGCTAATACCGCATGAAGTCGCGAGACCAAAG
TGGGGGACCTTCGGGCCTCACGCCTTCGGATGAGCCCAGATGAGATTAGCTGGTAGGTAAGGTAAAGGCTTACCTAGGCGACGATCTCTAGCGGGTCT
GAGAGGATAGCCCGCCACACTGGAACTGAGACACGGTCCAGACTCCTACGGGAGGCAGCAGTGGGGAATATTGCACAATGGGCGAAAGCCTGATGCAG
CCATGCCACGTGTGTGAAGAAGGCCTTCGGGTTGTAAAGCACTTTCAGCGAGGAGGAAGCGATAAATGCGAATACCATTTATTTTTGACGTTACTCGCAG
AAGAAGCACCGGCTAACTCCGTGCCAGCAGCCGCGGTAATACGGAGGGTGCGAGCGTTAATCGGAATAACTGGGCGTAAAGGGCATGTAGGTGGTGAG
TTAAGTCAGATGTGAAATCCCCGAGCTCAACTTGGGAATGGCATTTGAAACTGGGTCGCTAGGGTTTTCTAGGGGGGGTAGAATTCCAGGTGTAGCGGT
GAAATGCGTAGATATCTGGAGGAATACCGGTGGCGAAGGCGGCCCCCTGGAGAAAGACTGACGCTGAGGTGCGAAAGCGTGGGGAGCAAACAGGATTA
GATACCCTGGTAGTCCACGCTGTAAACGATGTCGATTTGGAGGTTGCGGTCTTGAACTGTGGCG

**> Seq11 [organism=Candidatus *Fukatsuia symbiotica*] 16S ribosomal RNA gene [host=*Acyrthosiphon pisum* isolate N116]**
AGAGTTTGATCCTGGCTCAGATTGAACGCTGGCGGCAGGCCTAACACATGCAAGTCGAGCGGCATCGGGAAGGTAGTTTGCTATCTTTGCCGGCGAGCG
GCGGACGGGTGAGTAAAGTCTGGGGATCTGCCTGATGGAGGGGGATAACTACTGGAAACGGTAGCTAATACCGCATGATGTTACGCGACCAAAGCGGG
GGACCTCCGGGCCTCGCGCCATCAGATGAACCCAGATGGGTTTAGCTAGTAGGAGAGGTAATGGCTCCCCTAGGCGACGATCCCTAGCTGGTCTGAGA
GGATAACCAGCCACACTGGAACTGAGAGACGGTCCAGACTCCTACGGGAGGCAGCAGTGGGGAATATTGCACAATGGGCGCAAGCCTGATGCAGCCAT
GCCGCGTGTGTGAAGAAGGCCTTCGGGTTGTAAAGCACTTTCAGCGAGGAGGAATGAAGCAATGCAAAGAGTGTTGCTAATGGACGTTACTCGCAGAAG
AAGCACCGGCTAACTCCGTGCCAGCAGCCGCGGTAATACGGAGGGTGCGAGCGTTAATCGGAATTACTGGGCGTAAAGAGCACGTAGGCGGTTTCTTAA
GTCAGACGTGAAATCCCCGAGCTTCACTTGGGAACGGCATTTGAAACTGAGAGTCTAGAGTTTTGTAGAGGGGGGTAGAATTCCAGGTGTAGCGGTGAA
ATGCGTAGATATCTGGAGGAATACCGGTGGCGAAGGCGGCCCCCTGGACAGAGACTGACGCTGAGGTGCGAAAGCGTGGGTAGCAAACAGGATTAGAT
ACCCTGGTAGTCCACGCTGTAAACGATGTCGATTTGTA

**> Seq12 [organism=Candidatus *Hamiltonella defensa*] 16S ribosomal RNA gene [host=*Acyrthosiphon pisum* isolate N116]**
AGAGTTTGATCCTGGCTCAGATTGAACACTGGTGGCAGGCCTAACACATACAAGTCGAGCGGCATCGAGTGAGCGCAGTTTACTGAGTTCATGTCGGCG
AGCGGCGGACGGGTGAGTAAAGTCTGGGAATCTGGCCGAAGGAGGGGGATAACTGCTGGAAACGGCAGCTAATACCGCATGAAGTCGCGAGACCAAAG
TGGGGGACCTTCGGGCCTCACGCCTTCGGATGAGCCCAGATGAGATTAGCTGGTAGGTAAGGTAAAGGCTTACCTAGGCGACGATCTCTAGCGGGTCT
GAGAGGATAGCCCGCCACACTGGAACTGAGACACGGTCCAGACTCCTACGGGAGGCAGCAGTGGGGAATATTGCACAATGGGCGAAAGCCTGATGCAG
CCATGCCACGTGTGTGAAGAAGGCCTTCGGGTTGTAAAGCACTTTCAGCGAGGAGGAAGCGATAAATGCGAATACCATTTATTTTTGACGTTACTCGCAG
AAGAAGCACCGGCTAACTCCGTGCCAGCAGCCGCGGTAATACGGAGGGTGTGAGCGTTAATCGGAATAACTGGGCGTAAAGGGCATGTAGGCGGTGAG
TTAAGTCAGATGTGAAATCCCCGAGCTCAACTTGGGAATGGCATTTGAAACTGGGTCGCTAGAGTTTTCTAGAGGGGGGTAGAATTCCAGGTGTAGCGGT
GAAATGCGTAGATATCTGGAGGAATACCGGTGGCGAAGGCGGCCCCCTGGAGAAAGACTGACGCTGAGGTGCGAAAGC

**> Seq13 [organism=Candidatus *Hamiltonella defensa*] 16S ribosomal RNA gene [host=*Acyrthosiphon pisum* isolate N116]**
AGAGTTTGATCCTGGCTCAGATTGAACACTGGTGGCAGGCCTAACACATGCAAGTCGAGCGGCATCGAGTGAGCGCAGTTTACTGAGTTCATGTCGGCG
AGCGGCGGACGGGTGAGTAAAGTCTGGGAATCTGGCCGAAGGAGGGGGATAACTGCTGGAAACGGCAGCTAATACCGCATGAAGTCGCGAGACCAAAG
TGGGGGACCTTCGGGCCTCACGCCTTCGGATGAGCCCAGATGAGATTAGCTGGTAGGTAAGGTAAGGGCTTACCTAGGCGACGATCTCTAGCGGGTCT
GAGAGGATAGCCCGCCACACTGGAACTGAGACACGGTCCAGACTCCTACGGGAGGCAGCAGTGGGGAATATTGCACAATGGGCGAAAGCCTGATGCAG
CCATGCCACGTGTGTGAAGAAGGCCTTCGGGTTGTAAAGCACTTTCAGCGAGGAGGAAGCGATAAATGCGAATACCATTTATTTTTGACGTTACTCGCAG
AAGAAGCACCGGCTAACTCCGTGCCAGCAGCCGCGGTAATACGGAGGGTGCGAGCGTTAATCGGAATAACTGGGCGTAAAGGGCATGTAGGCGGTGAG
TTAAGTCAGATGTGAAATCCCCGAGCTCAACTTGGGAATGGCATTTGAAACTGGGTCGCTAGAGTTTTCTAGAGGGGGGTAGAATTCCAGGTGTAGCGGT
GAAATGCGTAGATATCTGGAGGAATACCGGTGGCGAAGGCGGCCCCCTGGAGAAAGACTGACGCTGAGGTGCGAAAGCG

**> Seq14 [organism=Candidatus *Fukatsuia symbiotica*] 16S ribosomal RNA gene [host=*Acyrthosiphon pisum* isolate N116]**
GGTTACCTTGTTACGACTTCACCCCAGTCATGGTTCACAAAGTGGTAAGCGCCATCCCAAAGGTTAAGCTACCTACTTCTTTTGCAAAACACTCCCATGGT
GTGACGGGCGGTGTGTACAAGGCCCGGGAACGTATTCACCGTAGCATTCTGATCCACGATTACTAGCGATTCCGACTTCATGGAGTCGAGTTGCAGACT
CCAATCCGGACTACGACGTACTTTATGAGGTCCGCTCACCCTCGCAGGCTCGCTTCTCTTTGTATACGCCATTGTAGCACGTGTGTAGCCCTACTCGTAA
GGGCCATGATGACTTGACGTCATCCCCACCTTCCTCCGGTTTATCACCGGCAGTCTCTCTTGAGTTCCCACCTCTACGTGCTGGCAACAAAAGATAAGGG
TTGCGCTCGTTGCGGGACTTAACCCAACATTTCACAACACGAGCTGACGACAGCCATGCAGCACCTGTCTCAAAGCTCCCCGAAGGGCACGTCAACATC
TCTGTCGACTCCTCTGGATGTCAAGAGTAGGTAAGGTTCTTCGCGTTGCATCGAATTAAACCACATGCTCCACCGCTTGTGCGGGCCCCCGTCAATTCAT
TTGAGTTTTAACCTTGCGGCCGTACTCCCCAGGCGGTCGATTTAACGCGTTTGCTCCGCAAGCCACAGTTTATAACCACAACCTACAAATCGACATCGTTT
ACAGCGTGGACTACCAGGGTATCTAATCCTGTTTGCTACCCTCGCTTTCGCACCTCAGCGTCAGTCTCTGTCCAGGGGGCCGCCTTCGCCACCGGTATT
CCTCCAGATATCTACGCATTTCACCGCTACACCTGGAAATTCTACCCCCCCTCTACAAAACTCT

**> Seq15 [organism= Candidatus *Serratia symbiotica*] 16S ribosomal RNA gene [host=*Acyrthosiphon pisum* isolate N116]**
AGAGTTTGATCCTGGCTCAGATTGAACGCTGGCGGCAGGCCTAACACATGCAAGTCGAGCGGTAGCACAAGAGAGCTTGCTCTCTGGGTGACGAGCGG
CGGACGGGTGAGTAATGTCTGGGAAACTGCCTGATGGCGGGGGATAACTAGTGGAAACGGTAGCTAATACCGCATAACGTCGCAAGACCAAAGTGGGG
GACCTTCGGGCCTCACGCCATCAGATGTGCCCAGGTGGGATTAGCTGGTAGGTGGGGTAACGGCTCACCTAGGCGACGATCCCTAGCTGGTCTGAGAG
GATGACCAGCCACACTGGAACTGAGACACGGTCCAGACTCCTACGGGAGGCAGCAGTGGGGAATATTGCACAATGGGCGCAAGCCTGATGCAGCCATG
CCGCGTGTGTGAAGAAGGCCTTCGGGTTGTAAAGCACTTTCAGCGAGGAGAAAGGGTAATGTGTTAATAAGACATTGCATTGACGTTACTCGCAGAAGAA
GCACCGGCTAACTCCGTGCCAGCAGCCGCGGTAATACGGAGGGTGCAAGCGTTAATCGGAATTACTGGGCGTAAAGCGCACGCAGGCGGTTTGTTAAG
TCAGATGTGAAATCCCCGCGCTCAACGTAGGAACGGCATTTGAGACTGGCAAGCTAGAGTCTTGTAGAGGGGGGTAGAATTCCAGGTGTAGCGGTGAAA
TGCGTAGAGATCTGGAGGAATACCGGTGGCGAAGGCGGCCCCCTGGACAAAGACTGACGCTCAGGTGCGAAAGC

**> Seq16 [organism=Candidatus *Fukatsuia symbiotica*] 16S ribosomal RNA gene [host=*Acyrthosiphon pisum* isolate N116]**
GGTTACCTTGTTACGACTTCACCCCAGTCATGGTTCACAAAGTGGTAAGCGCCATCCCAAAGGTTAAGCTACCTACTTCTTTTGCAAAACACTCCCATGGT
GTGACGGGCGGTGTGTACAAGGCCCGGGAACGTATTCACCGTAGCATTCTGATCCACGATTACTAGCGATTCCGACTTCATGGAGTCGAGTTGCAGACT
CCAATCCGGACTACGACGTACTTTATGAGGTCCGCTCACCCTCGCAGGCTCGCTTCTCTTTGTATACGCCATTGTAGCACGTGTGTAGCCCTACTCGTAA
GGGCCATGATGACTTGACGTCATCCCCACCTTCCTCCGGTTTATCACCGGCAGTCTCTCTTGAGTTCCCACCTCTACGTGCTGGCAACAAAAGATAAGGG
TTGCGCTCGTTGCGGGACTTAACCCAACATTTCACAACACGAGCTGACGACAGCCATGCAGCACCTGTCTCAAAGCTCCCCGAAGGGCACGTCAACATC
TCTGTCGACTCCTCTGGATGTCAAGAGTAGGTAAGGTTCTTCGCGTTGCATCGAATTAAACCACATGCTCCACCGCTTGTGCGGGCCCCCGTCAATTCAT
TTGAGTTTTAACCTTGCGGCCGTACTCCCCAGGCGGTCGATTTAACGCGTTTGCTCCGCAAGCCACAGTTTATAACCACAACCTACAAATCGACATCGTTT
ACAGCGTGGACTACCAGGGTATCTAATCCTGTTTGCTACCCACGCTTTCGCACCTCAGCGTCAGTCTCTGTCCAGGGGGG

**> Seq17 [organism=Candidatus *Hamiltonella defensa*] 16S ribosomal RNA gene [host=*Acyrthosiphon pisum* isolate N116]**
AGAGTTTGATCCTGGCTCAGATTGAACACTGGTGGCAGGCCTAACACATGCAAGTCGAGCGGCATCGAGTGAGCGCAGTTTACTGAGTTCATGTCGGCG
AGCGGCGGACGGGTGAGTAAAGTCTGGGAATCTGGCCGAAGGAGGGGGATAACTGCTGGAAACGGCAGCTAATACCGCATGAAGTCGCGAGACCAAAG
TGGGGGACCTTCGGGCCTCACGCCTTCGGATGAGCCCAGATGAGATTAGCTGGTAGGTAAGGTAAAGGCTTACCTAGGCGACGATCTCTAGCGGGTCT
GAGAGGATAGCCCGCCACACTGGAACTGAGACACGGTCCAGACTCCTACGGGAGGCAGCAGTGGGGAATATTGCACAATGGGCGAAAGCCTGATGCAG
CCATGCCACGTGTGTGAAGAAGGCCTTCGGGTTGTAAAGCACTTTCAGCGAGGAGGAAGCGATAAATGCGAATACCATTTATTTTTGACGTTACTCGCAG
AAGAAGCACCGGCTAACTCCGTGCCAGCAGCCGCGGTAATACGGAGGGTGCGAGCGTTAATCGGAATAACTGGGCGTAAAGGGCATGTAGGCGGTGAG
TTAAGTCAGATGTGAAATCCCCGAGCTCAACTTGGGAATGGCATTTGAAACTGGGTCGCTAGAGTTTTCTAGAGGGGGGTAGAATTCCAGGTGTAGCGGT
GAAATGCGTAGATATCTGGAGGAATACCGGTGGCGAAGGCGGCCCCCTGGAGAAAGACTGACGCTGAGGTGCGAAAGC

**> Seq18 [organism=Candidatus *Fukatsuia symbiotica*] 16S ribosomal RNA gene [host=*Acyrthosiphon pisum* isolate N116]**
AGAGTTTGATCCTGGCTCAGATTGAACGCTGGCGGCAGGCCTAACACATGCAAGTCGAACGGCATCGGGAAGGTAGCTTGCTATCTTTGCCGGCGAGCG
GCGGACGGGTGAGTAAAGTCTGGGGATCTGCCTGATGGAGGGGGATAACTACTGGAAACGGTAGCTAATACCGCATGATGTTACGCGACCAAAGCGGG
GGACCTCCGGGCCTCGCGCCATCAGATGAACCCAGATGGGATTAGCTAGTAGGAGAGGTAATGGTTCCCCTAGGCGACGATCCCTAGCTGGTCTGAGA
GGATAACCAGCCACACTGGAACTGAGAGACGGTCCAGACTCCTACGGGAGGCAGCAGTGGGGAATATTGCACAATGGGCGCAAGCCTGATGCAGCCAT
GCCGCGTGTGTGAAGAAGGCCTTCGGGTTGTAAAGCACTTTCAGCGAGGAGGAATGAAGCAATGCAAAGAGTGTTGCTAATGGACGTTACTCGCAGAAG
AAGCACCGGCTAACTCCGTGCCAGCAGCCGCGGTAATACGGAGGGTGCGAGCGTTAATCGGAATTACTGGGCGTAAAGGGCACGTAGGCGGTTTCTTA
AGTCAGATGTGAAATCCCCGAGCTTCACTTGGGAACGGCATTTGAAACTGAGAGTCTAGAGTTTTGTAGAGGGGGGTAGAATTCCAGGTGTAGCGGTGAA
ATGCGT

**> Seq19 [organism=Candidatus *Hamiltonella defensa*] 16S ribosomal RNA gene [host=*Acyrthosiphon pisum* isolate N116]**
AGAGTTTGATCCTGGCTCAGATTGAACACTGGTGGCAGGCCTAACACATGCAAGTCGAGCGGCATCGAGTGAGCGCAGTTTACTGAGTTCATGTCGGCG
AGCGGCGGACGGGTGAGTAAAGTCTGGGAATCTGGCCGAAGGAGGGGGATAACTGCTGGAAACGGCAGCTAATACCGCATGAAGTCGCGAGACCAAAG
TGGGGGACCTTCGGGCCTCACGCCTTCGGATGAGCCCAGATGAGATTAGCTGGTAGGTAAGGTAAAGGCTTACCTAGGCGACGATCTCTAGCGGGTCT
GAGAGGATAGCCCGCCACACTGGAACTGAGACACGGTCCAGACTCCTACGGGAGGCAGCAGTGGGGAATATTGCACAATGGGCGAAAGCCTGATGCAG
CCATGCCACGTGTGTGAAGAAGGCCTTCGGGTTGTAAAGCACTTTCAGCGAGGAGGAAGCGATAAATGCGAATACCATTTATTTTTGACGTTACTCGCAG
AAGAAGCACCGGCTAACTCCGTGCCAGCAGCCGCGGTAATACGGAGGGTGCGAGCGTTAATCGGAATAACTGGGCGTAAAGGGCATGTAGGCGGTGAG
TTAAGTCAGATGTGAAATCCCCGAGCTCAACTTGGGAATGGCATTTGAAACTGGGTCGCTAGAGTTTTCTAGAGGGGGGTAGAATTCCAGGTGTAGCGGT
GAAATGCGTAGATATCTGGAGGAA

**> Seq20 [organism=*Buchnera aphidicola*] 16S ribosomal RNA gene [host=*Acyrthosiphon pisum* isolate N116]**
GGTTACCTTGTTACGACTTCACCCCAGTCATGAATCACAAAGTGGTAAGCGCCTTCCTTTTAAAGGGTTAGGATACCTGCTTCTTTTGCAACCCACTCCCA
TGGTGTGACGGGCGGTGTGTACAAGGCCCGGGAACGTATTCACCGTGGCATTCTGATCCACGATTACTAGCGATTCCGACTTCGTGGAGTCGAGTTGCA
GACTCCatTCCGGACTACGATTTACTTTATGAGGTTTGCTTGTCTTTGCAGATTTGCTTCTCTTTGTATAAACCATTGTAGCACGTGTGTAGCCCTGGTCGTA
AGGGCCATGATGACTTGACGTCGTCCCCACCTTCCTCCGGTTTATAACCGGCAGTCTCCTCTGAGTTCCCGGCCGAACCGCTGGCAACAGGGGATAAGG
GTTGCGCTCGTTGCGGGACTTAACCCAACATTTCACAACACGAGCTGACGACAGCCATGCAGCACCTGTCTCACAGCTCCCGAAGGCACTTCTTTATTTC
TAAAGAATTCTGTGGATGTCAAGACCAGGTAAGGTTTTTCGCGTTGCATCGAATTAAACCACATGCTCCACCGCTTGTGCGGGCCCCCGTCAATTCATTTG
AGTTTTAGCCTTGCGGCCGTACTCCCCAGGCGGTCGACTTAATGCGTTAGCTTCGGAAGTCACTTCTCTTGGAAACAACCTCCAAGTCGACATCGTTTAC
GGCATGGACTACCAGGGTATCTAATCCTGtTTGCTCCCCACGCtTTCGCGCCTCAGTGTC

**> Seq21 [organism=Candidatus *Hamiltonella defensa*] 16S ribosomal RNA gene [host=*Acyrthosiphon pisum* isolate N116]**
AGAGTTTGATCCTGGCTCAGATTGAACACTGGTGGCAGGCCTAACACATGCAAGTCGAGCGGCATCGAGTGAGCGCAGTTTACTGAGTTCATGTCGGCG
AGCGGCGGACGGGTGAGTAAAGTCTGGGAATCTGGCCGAAGGAGGGGGATAACTGCTGGAAACGGCAGCTAATACCGCATGAAGTCGCGAGACCAGA
GTGGGGGACCTTTGGGCCTCACGCCTTCGGATGAGCCCAGATGTGATTAGCTGGTAGGTAAGGTAAAGGCTTACCTAGGCGACGATCTCTAGCGGGTCT
GAGAGGATAGCCCGCCACACTGGAACTGAGACACGGTCCAGACTCCTACGGGAGGCAGCAGTGGGGAATATTGCCCAATGGGCGAAAGCCTGATGCAG
CCATGCCACGTGTGTGAAGAAGGCCTTCGGGTTGTAAAGCACTTTCAGCGAGGAGGAAGCGATAAATGCGAATACCATTTATTTTTGACGTTACTCGCAG
AAGAAGCACCGGCTAACTCCGTGCCAGCAGCCGCGGTAATACGGAGGGTGCGAGCGTTAATCGGAATAACTGGGCGTAAAGGGCATGTAGGCGGTGAG
TTAAGCCAGATGTGAAATCCCCGAGCTCAACTTGGGAATGGCATTTGAAACTGGGTCGCTAGAGTTTTCTAGAGGGGGGTAGAATTCCAGGTGTAGCGGT

**> Seq22 [organism=Candidatus *Hamiltonella defensa*] 16S ribosomal RNA gene [host=*Acyrthosiphon pisum* isolate N116]**
AGAGTTTGATCCTGGCTCAGATTGAACACTGGTGGCAGGCCTAACACATGCAAGTCGAGCGGCATCGAGTGAGCGCAGTTTACTGAGTTCATGTCGGCG
AGCGGCGGACGGGTGAGTAAAGTCTGGGAATCTGGCCGAAGGAGGGGGATAACTGCTGGAAACGGCAGCTAATACCGCATGAAGTCGCGAGACCAAAG
TGGGGGACCTTCGGGCCTCACGCCTTCGGATGAGCCCAGATGAGATTAGCTGGTAGGTAAGGTAAAGGCTTACCTAGGCGACGATCTCTAGCGGGTCT
GAGAGGATAGCCCGCCACACTGGAACTGAGACACGGTCCAGACTCCTACGGGAGGCAGCAGTGGGGAATATTGCACAATGGGCGAAAGCCTGATGCAG
CCATGCCACGTGTGTGAAGAAGGCCTTCGGGTTGTAAAGCACTTTCAGCGAGGAGGAAGCGATAAATGCGAATACCATTTATTTTTGACGTTACTCGCAG
AAGAAGCACCGGCTAACTCCGTGCCAGCAGCCGCGGTAATACGGAGGGTGCGAGCGTTAATCGGAATAACTGGGCGTAAAGGGCATGTAGGCGGTGAG
TTAAGTCAGATGTGAAATCCCCGAGCTCAACTTGGGAATGGCATTTGAAACTGGGTCGCTAGAGTTTTCTAGAGGGGGGTAGAATTCCAGGTGTAGCGGT
GAAATGCGTAGATATCTGGAGGAATACCGGTGGCGAAGGCGGCCCCCTGGAGAAAGACTGACGCTGAGG

**> Seq23 [organism=Candidatus *Hamiltonella defensa*] 16S ribosomal RNA gene [host=*Acyrthosiphon pisum* isolate N116]**
GGTTACCTTGTTACGACTTCACCCCAGTCATGAATCACAAAGTGGTAAGCGCCCTCCTTGCGGTTTAGCTACCTACTTCTTTTGCAACCCACTCCCATGGT
GTGACGGGCGGTGTGTACAAGGCCCGGGAACGTATTCACCGTAGCATTCTGATCTACGATTACTAGCGATTCCGACTTCATGGAGTCGAGTTGCAGACT
CCAATCCGGACTACGACATACTTTCTGAGTTCCGCTTTCCCTCGCAGGTTCGCATCCCTTTGTATACGCCATTGTAGCACGTGTGTAGCCCTACTCGTAAG
GGCCATGATGACTTGACGTCGTCCCCACCTTCCTCCGGTTTATCACCGGCAGTCTCCTTTGAGTTCCCGCCTCTACGCGCTGGCAACAAAGGACAAGGG
ATGCGCTCGTTGCGGGACTTAACCCAACATTTCACAACACGAGCTGACGACAGCCATGCAGCACCTGTCTCACGGTTCCCGAAGGCACTTGCGCATCTC
TGCACAATTCCGTGGATGTCAAGAGTAGGTAAGGTTCTTCGCGTTGCATCGAATTAAACCACATGCTCCACCGCTTGTGCGGGCCCCCGTCAATTCATTT
GAGTTTTAACCTTGCGGCCGTACTCCCCAGGCGGTCGATTTAACGCGTTAGCTCCGGACGCCACAGTTCAAGACCGCAACCTCCAAATCGACATCGTTTA
CAGCGTGGACTACCAGGGTATCTAATCCTGTTTGCTCCCCACGCTTTCGCACCTCAGCGTCAGTCTTTCTCCAGGGGGCCGCCTTCGCCACCGGTATTC
CTCCAGATATCTACGCATTTCACCGCTACACCT

**> Seq24 [organism=Candidatus *Fukatsuia symbiotica*] 16S ribosomal RNA gene [host=*Acyrthosiphon pisum* isolate N116]**
AGAGTTTGATCCTGGCTCAGATTGAACGCTGGCGGCAGGCCTAACACATGCAAGTCGAGCGGCATCGGGAAGGTAGCTTGCTATCTTTGCCGGCGAGC
GGCGGACGGGTGAGTAAAGTCTGGGGATCTGCCTGATGGAGGGGGATAACTACTGGAAACGGTAGCTAATACCGCATGATGTTACGCGACCAAAGCGG
GGGACCTCCGGGCCTCGCGCCATCAGATGAACCCAGATGGGATTAGCTAGTAGGAGAGGTAATGGCTCCCCTAGGCGACGATCCCTAGCTGGTCTGAG
AGGATAACCAGCCACACTGGAACTGAGAGACGGTCCAGACTCCTACGGGAGGCAGCAGTGGGGAATATTGCACAATGGGCGCAAGCCTGATGCAGCCA
TGCCGCGTGTGTGAAGAAGGCCTTCGGGTTGTAAAGCACTTTCAGCGAGGAGGAATGAAGCAATGCAAAGAGTGTTGCTAATGGACGTTACTCGCAGAA
GAAGCACCGGCTAACTCCGTGCCAGCAGCCGCGGTAATACGGAGGGTGCGAGCGTTAATCGGAATTACTGGGCGTAAAGGGCACGTAGGCGGTTTCTT
AAGTCAGATGTGAAATCCCCGAGCTTCACTTGGGAACGGCATTTGAAACTGAGAGTCTAGAGTTTTGTAGAGGGGGGTAGAAT

**> Seq25 [organism=Candidatus *Hamiltonella defensa*] 16S ribosomal RNA gene [host=*Acyrthosiphon pisum* isolate N116]**
AGAGTTTGATCCTGGCTCAGATTGAACACTGGTGGCAGGCCTAACACATGCAAGTCGAGCGGCATCGAGTGAGCGCAGTTTACTGAGTTCATGTCGGCG
AGCGGCGGACGGGTGAGTAAAGTCTGGGAATCTGGCCGAAGGAGGGGGATAACTGCTGGAAACGGCAGCTAATACCGCATGAAGTCGCGAGACCAAAG
TGGGGGACCTTCGGGCCTCACGCCTTCGGATGAGCCCAGATGAGATTAGCTGGTAGGTAAGGTAAAGGCTTACCTAGGCGACGATCTCTAGCGGGTCT
GAGAGGATAGCCCGCCACACTGGAACTGAGACACGGTCCAGACTCCTACGGGAGGCAGCAGTGGGGAATATTGCACAATGGGCGAAAGCCTGATGCAG
CCATGCCACGTGTGTGAAGAAGGCCTTCGGGTTGTAAAGCACTTTCAGCGAGGAGGAAGCGATAAATGCGAATACCATTTATTTTTGACGTTACTCGCAG
AAGAAGCACCGGCTAACTCCGTGCCAGCAGCCGCGGTAATACGGAGGGTGCGAGCGTTAATCGGAATAACTGGGCGTAAAGGGCATGTAGGCGGTGAG
TTAAGTCAGATGTGAAATCCCCGAGCTCAACTTGGGAATGGCATTTGAAACTGGGTCGCTAGAGTTTTCTAGAGGGGGGTAGAATTCCAGGTGT

**Q1 aphid symbiont BLAST analysis**

Trimmed ‘Q1 clone *Acyrthosiphon pisum* symbiont 16s rRNA gene sequences used in the blast analysis shown in FASTA format. GenBank Accession numbers: MW971996 to MW972018

**> Seq1 [organism= Candidatus *Serratia symbiotica*] 16S ribosomal RNA gene [host=*Acyrthosiphon pisum* isolate Q1]**
AGAGTTTGATCCTGGCTCAGATTGAACGCTGGCGGCAGGCCTAACACATGCAAGTCGAGCGGTAGCACAAGAGAGCTTGCTCTCTGGGTGACGAGCGG
CGGACGGGTGAGTAATGTCTGGGAAACTGCCTGATGGCGGGGGATAACTAGTGGAAACGGTAGCTAATACCGCATAACGTCGCAAGACCAAAGTGGGG
GACCTTCGGGCCTCACGCCATCAGATGTGCCCAGGTGGGATTAGCTGGTAGGTGGGGTAACGGCTCACCTAGGCGACGATCCCTAGCTGGTCTGAGAG

GATGACCAGCCACACTGGAACTGAGACACGGTCCAGACTCCTACGGGAGGCAGCAGTGGGGAATATTGCACAATGGGCGCAAGCCTGATGCAGCCATG
CCGCGTGTGTGAAGAAGGCCTTCGGGTTGTAAAGCACTTTCAGCGAGGAGAAAGGGTAATGTGTTAATAAGACATTGCATTGACGTTACTCGCAGAAGAA
GCACCGGCTAACTCCGTGCCAGCAGCCGCGGTAATACGGAGGGTGCAAGCGTTAATCGGAATTACTGGGCGTAAAGCGCACGCAGGCGGTTTGTTAAG
TCAGATGTGAAATCCCCGCGCTCAACGTAGGAACGGCATTTGAGACTGGCAAGCTAGAGTCTTGTAGAGGGGG

**> Seq2 [organism= Candidatus *Serratia symbiotica*] 16S ribosomal RNA gene [host=*Acyrthosiphon pisum* isolate Q1]**
AGAGTTTGATCCTGGCTCAGATTGAACGCTGGCGGCAGGCCTAACACATGCAAGTCGAGCGGTAGCACAAGAGAGCTTGCTCTCTGGGTGACGAGCGG
CGGACGGGTGAGTAATGTCTGGGAAACTGCCTGATGGCGGGGGATAACTAGTGGAAACGGTAGCTAATACCGCATAACGTCGCAAGACCAAAGTGGGG
GACCTTCGGGCCTCACGCCATCAGATGTGCCCAGGTAGGATTAGCTGGTAGGTGGGGTAACGGCTCACCTAGGCGACGATCCCTAGCTGGCCTGAGAG
GATGACCAGCCACACTGGAACTGAGACACGGTCCAGACTCCTACGGGAGGCAGCAGTGGGGAATATTGCACAATGGGCGCAAGCCTGATGCAGCCATG
CCGCGTGTGTGAAGAGGGCCTTCGGGTTGTAAAGCACTTTCAGCGAGGAGAAAGGGTAATGTGTTAATAAGACATTGCATTGACGTTACTCGCAGAAGAA
GCACCGGCTAACTCCGTGCCTGCAGCCGCGGTAATACGGAGGGTGCAAGCGTTAATCGGAATTACTGGGCGTAAAGCGCACGCAGGCGGTTTGTTAAG
TCAGATGTGAAATCCCCGCGCTCAACGTGGGAACGGCATTTGAGACTGGCAAGCTAGAGTCTTGTAGAGGGGGGTAGAATTC

**> Seq3 [organism= Candidatus *Serratia symbiotica*] 16S ribosomal RNA gene [host=*Acyrthosiphon pisum* isolate Q1]**
CGGCAGGCCTAACACATGCAAGTCGAGCGGTAGCACAAGAGAGCTTGCTCTCTGGGTGACGAGCGGCGGACGGGTGAGTAATGTCTGGGAAACTGCCT
GATGGCGGGGGATAACTAGTGGAAACGGTAGCTAATACCGCATAACGTCGCAAGACCAAAGTGGGGGACCTTCGGGCCTCACGCCATCAGATGTGCCC
AGGTGGGATTAGCTGGTAGGTGGGGTAACGGCTCACCTAGGCGACGATCCCTAGCTGGTCTGAGAGGATGACCAGCCACACTGGAACTGAGACACGGT
CCAGACTCCTACGGGAGGCAGCAGTGGGGAATATTGCACAATGGGCGCAAGCCTGATGCAGCCATGCCGCGTGTGTGAAGAAGGCCTTCGGGTTGTAA
AGCAC

**> Seq4 [organism= Candidatus *Serratia symbiotica*] 16S ribosomal RNA gene [host=*Acyrthosiphon pisum* isolate Q1]**
AGAGTTTGATCCTGGCTCAGATTGAACGCTGGCGGCAGGCCTAACACATGCAAGTCGAGCGGTAGCACAAGAGAGCTTGCTCTCTGGGTGACGAGCGG
CGGACGGGTGAGTAATGTCTGGGAAACTGCCTGATGGCGGGGGATAACTAGTGGAAACGGTAGCTAATACCGCATAACGTCGCAAGACCAAAGTGGGG
GACCTTCGGGCCTCACGCCATCAGATGTGCCCAGGTAGGATTAGCTGGTAGGTGGGGTAACGGCTCACCTAGGCGACGATCCCTAGCTGGTCTGAGAG
GATGACCAGCCACACTGGAACTGAGACACGGTCCAGACTCCTACGGGAGGCAGCAGTGGGGAATATTGCACAATGGGCGCAAGCCTGATGCAGCCATG
CCGCGTGTGTGAAGAAGGCCTTCGGGTTGTAAAGCACTTTCAGCGAGGAGAAAGGGTAATGTGTTAATAAGACATTGCATTGACGTTACTCGCAGAAGAA
GCACCGGCTAACTCCGTGCCAGCAGCCGCGGTAATACGGAGGGTGCAAGCGTTAATCGGAATTACTGGGCGTAAAGCGCACGCAGGCGGTTTGTTAAG
TCAGATGTGAAATCCCCGCGCTCAACGTGGGAACGGCATTTGAGACTGGCAAGCTAGAGTCTTGTAGAGGGGGGTAGAATT

**> Seq5 [organism= Candidatus *Serratia symbiotica*] 16S ribosomal RNA gene [host=*Acyrthosiphon pisum* isolate Q1]**
AGAGTTTGATCCTGGCTCAGATTGAACGCTGGCGGCAGGCCTAACACATGCAAGTCGAGCGGTAGCACAAGAGAGCTTGCTCTCTGGGTGACGAGCGG
CGGACGGGTGAGTAATGTCTGGGAAACTGCCTGATGGCGGGGGATAACTAGTGGAAACGGTAGCTAATACCGCATAACGTCGCAAGACCAAAGTGGGG
GACCTTCGGGCCTCACGCCATCAGATGTGCCCAGGTGGGATTAGCTGGTAGGTGGGGTAACGGCTCACCTAGGCGACGATCCCTAGCTGGTCTGAGAG
GATGACCAGCCACACTGGAACTGAGACACGGTCCAGACTCCTACGGGAGGCAGCAGTGGGGAATATTGCACAATGGGCGCAAGCCTGATGCAGCCAGG
CCGCGTGTGTGAAGAAGGCCTTCGGGTTGTAAAGCACTTTCAGCGAGGAGAAAGGGTAATGTGTTAATAAGACATTGCATTGACGTTACTCGCAGAAGAA
GCACCGGCTAACTCCGTGCCAGCAGCCGCGGTAATACGGAGGGTGCAAGCGTTAATCGGAATTACTGGGCGTAAAGCGCACGCAGGCGGTTTGTTAAG
TCAGATGTGAAATCCCCGCGCTCAACGTAGGAACGGCATTTGAGACTGGCAA

**> Seq6 [organism= Candidatus *Serratia symbiotica*] 16S ribosomal RNA gene [host=*Acyrthosiphon pisum* isolate Q1]**
AGAGTTTGATCCTGGCTCAGATTGAACGCTGGCGGCAGGCCTAACACATGCAAGTCGAGCGGTAGCACAAGAGAGCTTGCTCTCTGGGTGACGAGCGG
CGGACGGGTGAGTAATGTCTGGGAAACTGCCTGATGGCGGGGGATAACTAGTGGAAACGGTAGCTAATACCGCATAACGTCGCAAGACCAAAGTGGGG
GACCTTCGGGCCTCACGCCATCAGATGTGCCCAGGTGGGATTAGCTGGTAGGTGGGGTAACGGCTCACCTAGGCGACGATCCCTAGCTGGTCTGAGAG
GATGACCAGCCACACTGGAACTGAGACACGGTCCAGACTCCTACGGGAGGCAGCAGTGGGGAATATTGCACAATGGGCGCAAGCCTGATGCAGCCATG
CCGCGTGTGTGAAGAAGGCCTTCGGGTTGTAAAGCACTTTCAGCGAGGAGAAAGGGTAATGTGTTAATAAGACATTGCATTGACGTTACTCGCAGAAGAA
GCACCGGCTAACTCCGTGCCAGCAGCCGCGGTAATACGGAGGGTGCAAGCGTTAATCGGAATTACTGGGCGTAAAGCGCACGCAGGCGGTTTGTTAAG
TCAGATGTGAAATCCCCGCGCTCAACGTGGGAACGGCATTTGAGACTGGCAAGCTAGAGTCTTGTAGAGGGGGGTAGAATTCCAGGTGTAGCGGTGAAA
TGCG

**> Seq7 [organism= Candidatus *Serratia symbiotica*] 16S ribosomal RNA gene [host=*Acyrthosiphon pisum* isolate Q1]**
AGAGTTTGATCCTGGCTCAGATTGAACGCTGGCGGCAGGCCTAACACATGCAAGTCGAGCGGTAGCACAAGAGAGCTTGCTCTCTGGGTGACGAGCGG
CGGACGGGTGAGTAATGTCTGGGAAACTGCCTGATGGCGGGGGATAACTAGTGGAAACGGTAGCTAATACCGCATAACGTCGCAAGACCAAAGTGGGG
GACCTTCGGGCCTCACGCCATCAGATGTGCCCAGGTGGGATTAGCTGGTAGGTGGGGTAACGGCTCACCTAGGCGACGATCCCTAGCTGGTCTGAGAG
GATGACCAGCCACACTGGAACTGAGACACGGTCCAGACTCCTACGGGAGGCAGCAGTGGGGAATATTGCACAATGGGCGCAAGCCTGATGCAGCCATG
CCGCGTGTGTGAAGAAGGCCTTCGGGTTGTAAAGCACTTTCAGCGAGGAGAAAGGGTAATGTGTTAATAAGACATTGCATTGACGTTACTCGCAGAAGAA
GCACCGGCTAACTCCGTGCCAGCAGCCGCGGTAATACGGAGG

**> Seq8 [organism= *Buchnera aphidicola*] 16S ribosomal RNA gene [host=*Acyrthosiphon pisum* isolate Q1]**
GGTTACCTTGTTACGACTTCACCCCAGTCATGAATCACAAAGTGGTAAGCGCCTTCCTTTTAAAGGGTTAGGATACCTGCTTCTTTTGCAACCCACTCCCA
TGGTGTGACGGGCGGTGTGTACAAGGCCCGGGAACGTATTCACCGTGGCATTCTGATCCACGATTACTAGCGATTCCGACTTCGTGGAGTCGAGTTGCA
GACTCCAGTCCGGACTACGATTTACTTTATGAGGTTTGCTTGTCTTTGCAGATTTGCTTCTCTTTGTATAAACCATTGTAGCACGTGTGTAGCCCTGGTCGT
AAGGGCCATGATGACTTGACGTCGTCCCCACCTTCCTCCGGTTTATAACCGGCAGTCTCCTCTGAGTTCCCGGCCGAACCGCTGGCAACAGGGGATAAG
GGTTGCGCTCGTTGCGGGACTTAACCCAACATTTCACAACACGAGCTGACGACAGCCATGCAGCACCTGTCTCACAGCTCCCGAAGGCACTTCTTTATTT
CTAAAGAATTCTGTGGATGTCAAGACCAGGTAAGGTTTTTCGCGTTGCATCGAATTAAACCACATGCTCCACCGCTTGTGCGGGCCCCCGTCAATTCATTT
GAGTTTTAGCCTTGCGGCCGTACTCCCCAGGCGGTCGACTTAATGCGTTAGCTTCGGAAGTCACTTCTCTTGGAAACAACCTCCAAGTCGACATCGTTTA
CGGCATGGACCACCAGGGTATCTAATCCTGTTTGCTCCCCACGCTTTCGCGCCTCAGTGTCAGTTTTT

**> Seq9 [organism= Candidatus *Serratia symbiotica*] 16S ribosomal RNA gene [host=*Acyrthosiphon pisum* isolate Q1]**
AGAGTTTGATCCTGGCTCAGATTGAACGCTGGCGGCAGGCCTAACACATGCAAGTCGAGCGGTAGCACAAGAGAGCTTGCTCTCTGGGTGACGAGCGG
CGGACGGGTGAGTAATGTCTGGGAAACTGCCTGATGGCGGGGGATAACTAGTGGAAACGGTAGCTAATACCGCATAACGTCGCAAGACCAAAGTGGGG
GACCTTCGGGCCTCACGCCATCAGATGTGCCCAGGTAGGATTAGCTGGTAGGTGGGGTAACGGCTCACCTAGGCGACGATCCCTAGCTGGTCTGAGAG
GATGACCAGCCACACTGGAACTGAGACACGGTCCAGACTCCTACGGGAGGCAGCAGTGGGGAATATTGCACAATGGGCGCAAGCCTGATGCAGCCATG
CCGCGTGTGTGAAGAAGGCCTTCGGGTTGTAAAGCACTTTCAGCGAGGAGAAAGGGTAATGTGTTAATAAGACATTGCATTGACGTTACTCGCAGAAGAA
GCACCGGCTAACTCCGTGCCAGCAGCCGCGGTAATACGGAGGGTGCAAGCGTTAATCGGAATTACTGGGCGTAAAGCGCACGCAGGCGGTTTGTTAAG
TCAGATGTGAAATCCCCGCGCTCAACGTGGGAACGGCATTTTGAGACTG

**> Seq10 [organism= Candidatus *Serratia symbiotica*] 16S ribosomal RNA gene [host=*Acyrthosiphon pisum* isolate Q1]**
AGAGTTTGATCCTGGCTCAGATTGAACGCTGGCGGCAGGCCTAACACATGCAAGTCGAGCGGTAGCACAAGAGAGCTTGCTCTCTGGGTGACGAGCAG
CGGACGGGTGAGTAATGTCTGGGAAACTGCCTGATGGCGGGGGATAACTAGTGGAAACGGTAGCTAATACCGCATAACGTCGCAAGACCAAAGTGGGG
GACCTTCGGGCCTCACGCCATCAGATGTGCCCAGGTGGGATTAGCTGGTAGGTGGGGTAACGGCTCACCTAGGCGACGATCCCTAGCTGGTCTGAGAG
GATGACCAGCCACACTGGAACTGAGACACGGTCCAGACTCCTACGGGAGGCAGCAGTGGGGAATATTGCACAATGGGCGCAAGCCTGATGCAGCCATG
CCGCGTGTGTGAAGAAGGCCTTCGGGTTGTAAAGCACTTTCAGCGAGGAGAAAGGGTAATGTGTTAATAAGACATTGCATTGACGTTACTCGCAGAAGAA
GCACCGGCTAACTCCGTGCCAGCAGCCGCGGTAATACGGAGGGTGCAAGCGTTAATCGGAATTACTGGGCGTAAAGCGCACGCAGGCGGTTTGTTAAG
TCAGATGTGAAATCCCCGCGCTCAACGTGGGAACGGCATTTGAGACTGGCAAGCTAGAGTCTTGTAGAGGGGGGTAGAATTCCAGG

**> Seq11 [organism= Candidatus *Serratia symbiotica*] 16S ribosomal RNA gene [host=*Acyrthosiphon pisum* isolate Q1]**
AGAGTTTGATCCTGGCTCAGATTGAACGCTGGCGGCAGGCCTAACACATGCAAGTCGAGCGGTAGCACAAGAGAGCTTGCTCTCTGGGTGACGAGCGG
CGGACGGGTGAGTAATGTCTGGGAAACTGCCTGATGGCGGGGGATAACTAGTGGAAACGGTAGCTAATACCGCATAACGTCGCAAGACCAAAGTGGGG
GACCTTCGGGCCTCACGCCATCAGATGTGCCCAGGTGGGATTAGCTGGTAGGTGGGGTAACGGCTCACCTAGGCGACGATCCCTAGCTGGTCTGAGAG
GATGACCAGCCACACTGGAACTGAGACACGGTCCAGACTCCTACGGGAGGCAGCAGTGGGGAATATTGCACAATGGGCGCAAGCCTGATGCAGCCATG
CCGCGTGTGTGAAGAAGGCCTTCGGGTTGTAAAGCACTTTCAGCGAGGAGAAAGGGTAATGTGTTAATAAGACATTGCATTGACGTTACTCGCAGAAGAA
GCACCGGCTAACTCCGTGCCAGCAGCCGCGGTAATACGGAGGGTGCAAGCGTTAATCGGAATTACTGGGCGTAAAGCGCACGCAGGCGGTTTGTTAAG
TCAGATGTGAAATCCCCGCGCTCAACGT

**> Seq12 [organism= Candidatus *Serratia symbiotica*] 16S ribosomal RNA gene [host=*Acyrthosiphon pisum* isolate Q1]**
AGAGTTTGATCCTGGCTCAGATTGAACGCTGGCGGCAGGCCTAACACATGCAAGTCGAGCGGTAGCACAAGAGAGCTTGCTCTCTGGGTGACGAGCGG
CGGACGGGTGAGTAATGTCTGGGAAACTGCCTGATGGCGGGGGATAACTAGTGGAAACGGTAGCTAATACCGCATAACGTCGCAAGACCAAAGTGGGG
GACCTTCGGGCCTCACGCCATCAGATGTGCCCAGGTGGGATTAGCTGGTAGGTGGGGTAACGGCTCACCTAGGCGACGATCCCTAGCTGGTCTGAGAG
GATGACCAGCCACACTGGAACTGAGACACGGTCCAGACTCCTACGGGAGGCAGCAGTGGGGAATATTGCACAATGGGCGCAAGCCTGATGCAGCCATG
CCGCGTGTGTGAAGAAGGCCTTCGGGTTGTAAAGCACTTTCAGCGAGGAGAAAGGGTAATGTGTTAATAAGACATTGCATTGACGTTACTCGCAGAAGAA
GCATCGGCTAACTCCGTGCCAGCAGCCGCGGTAATACGGAGGGTGCAAGCGTTAATCGGAATTACTGGGCGTAAAGCGCACGTAGGCGGTTTGTTAAGT
CAGATGTGAAATCCCCGCGCTCAACGTGGGAACGGCATTTGAGACTGGCAAGCTAGAGTCTTGTAGAGGGGGGTAGAATTCCAGGTGTAGCGGTGAAAT
GCGTAGAGATCTGGAGGAATACCGGTGGCGAAGGCGGCCCCCTGGACAAAGACTGACGCTCAGGTGCGAAAGCGTGGGGAGCAAACAGGATTAGATAC
CCTGGTAGTCCACGCTGTAAACGATGTCGATTTGGAGGTTGCGCCCTTGAGGGGTGGCTTCCGTAGCTAACGCGTTAAATCGACCGCCTGGGGAGTAC
G

**> Seq13 [organism= Candidatus *Serratia symbiotica*] 16S ribosomal RNA gene [host=*Acyrthosiphon pisum* isolate Q1]**
AGAGTTTGATCCTGGCTCAGATTGAACGCTGGCGGCAGGCCTAACACATGCAAGTCGAGCGGTAGCACAAGAGAGCTTGCTCTCTGGGTGACGAGCGG
CGGACGGGTGAGTAATGTCTGGGAAACTGCCTGATGGCGGGGGATAACTAGTGGAAACGGTAGCTAATACCGCATAACGTCGCAAGACCAAAGTGGGG
GACCTTCGGGCCTCACGCCATCAGATGTGCCCAGGTAGGATTAGCTGGTAGGTGGGGTAACGGCTCACCTAGGCGACGATCCCTAGCTGGTCTGAGAG
GATGACCAGCCACACTGGAACTGAGACACGGTCCAGACTCCTACGGGAGGCAGCAGTGGGGAATATTGCACAATGGGCGCAAGCCTGATGCAGCCATG
CCGCGTGTGTGAAGAAGGCCTTCGGGTTGTAAAGCACTTTCAGCGAGGAGAAAGGGTAATGTGTTAATAAGACATTGCATTGACGTTACTCGCAGAAGAA
GCACCGGCTAACTCCGTGCCAGCAGCCGCGGTAATACGGAGGGTGCAAGCGTTAATCGGAATTACTGGGCGTAAAGCGCACGCAGGCGGTTTGTTAAG
TCAGATGTGAAATCCCCGCGCTCAACGTGGGAACGGCATTTGAGACTGGCAAGCTAGAGTCTTGTAGAGGGGGGTAGAATTCCAGGTGTAGCGGTGAAA
TGCGTAGAGATCTGGAGGAATACCGGTGGCGAAGGCGGCCCCCTGGACAAAGACTGACGCTCAGGTGCGAAAGCGTGGGGAGCAAACAGGATTAGATA
CCCTGGTAGTCCACGCTGTAAACGATGTCGATTTGGAGGTTGCGCCCCTTG

**> Seq14 [organism= Candidatus *Serratia symbiotica*] 16S ribosomal RNA gene [host=*Acyrthosiphon pisum* isolate Q1]**
AGAGTTTGATCCTGGCTCAGATTGAACGCTGGCGGCAGGCCTAACACATGCAAGTCGAGCGGTAGCACAAGAGAGCCTGCTCTCTGGGTGACGAGCGG
CGGACGGGTGAGTAATGTCTGGGAAACTGCCTGATGGCGGGGGATAACTAGTGGAAACGGTAGCTAATACCGCATAACGTCGCAAGACCAAAGTGGGG
GACCTTCGGGCCTCACGCCATCAGATGTGCCCAGGTAGGATTAGCTGGTAGGTGGGGTAACGGCTCACCTAGGCGACGATCCCTAGCTGGTCTGAGAG
GATGACCAGCCACACTGGAACTGAGACACGGTCCAGACTCCTACGGGAGGCAGCAGTGGGGAATATTGCACAATGGGCGCAAGCCTGATGCAGCCATG
CCGCGTGTGTGAAGAAGGCCTTCGGGTTGTAAAGCACTTTCAGCGAGGAGAAAGGGTAATGTGTTAATAAGACATTGCATTGACGTTACTCGCAGAAGAA
GCACCGGCTAACTCCGTGCCAGCAGCCGCGGTAATACGGAGGGTGCAAGCGTTAATCGGAATTACTGGGCGTAAAGCGCACGCAGGCGGTTTGTTAAG
TCAGATGTGAAATCCCCGCGCTCAACGTGGGAACGGCATTTGAGACTGGCAAGCTAGAGTCTTGTAGAGGGGGGTAGAATTCCAGGTGTAGCGGTGAAA
TGCGTAGAGATCTGGAGGA

**> Seq15 [organism= Candidatus *Serratia symbiotica*] 16S ribosomal RNA gene [host=*Acyrthosiphon pisum* isolate Q1]**
AGAGTTTGATCCTGGCTCAGATTGAACGCTGGCGGCAGGCCTAACACATGCAAGTCGAGCGGTAGCACAAGAGAGCTTGCTCTCTGGGTGACGAGCGG
CGGACGGGTGAGTAATGTCTGGGAAACTGCCTGATGGCGGGGGATAACTAGTGGAAACGGTAGCTAATACCGCATAACGTCGCAAGACCAAAGTGGGG
GACCTTCGGGCCTCACGCCATCAGATGTGCCCAGGTGGGATTAGCTGGTAGGTGGGGTAACGGCTCACCTAGGCGACGATCCCTAGCCGGTCTGAGAG
GATGACCAGCCACACTGGAACTGAGACACGGTCCAGACTCCTACGGGAGGCAGCAGTGGGGAATATTGCACAATGGGCGCAAGCCTGATGCAGCCATG
CCGCGTGTGTGAAGAAGGCCTTCGGGTTGTAAAGCACTTTCAGCGAGGAGAAAGGGTAATGTGTTAATAAGACATTGCATTGACGTTACTCGCAGAAGAA
GCACCGGCTAACTCCGTGCCAGCAGCCGCGGTAATACAGAGGGTGCAAGCGTTAATCGGAATTACTGGGCGTAAAGCGCACGCAGGCGGTTTGTTAAG
TCAGATGTGAAATCCCCGCGCTCAACGTAGGAACGGCATTTGAGACTGGCAA

**> Seq16 [organism= Candidatus *Serratia symbiotica*] 16S ribosomal RNA gene [host=*Acyrthosiphon pisum* isolate Q1]**
AGAGTTTGATCCTGGCTCAGATTGAACGCTGGCGGCAGGCCTAACACATGCAAGTCGAGCGGTAGCACAAGAGAGCTTGCTCTCTGGGTGACGAGCGG
CGGACGGGTGAGTAATGTCTGGGAAACTGCCTGATGGCGGGGGATAACTAGTGGAAACGGTAGCTAATACCGCATAACGTCGCAAGACCAAAGTGGGG
GACCTTCGGGCCTCACGCCATCAGATGTGCCCAGGTGGGATTAGCTGGTAGGTGGGGTAACGGCTCACCTAGGCGACGATCCCTAGCTGGTCTGAGAG
GATGACCAGCCACACTGGAACTGAGACACGGTCCAGACTCCTACGGGAGGCAGCAGTGGGGAATATTGCACAAAGGGCGCAAGCCTGATGCAGCCATG
CCGCGTGTGTGAAGAAGGCCTTCGGGTTGTAAAGCACTTTCAGCGAGGAGAAAGGGTAATGTGTTAATAAGACATTGCATTGACGTTACTCGCA

**> Seq17 [organism= *Buchnera aphidicola*] 16S ribosomal RNA gene [host=*Acyrthosiphon pisum* isolate Q1]**
AGAGTTTGATCCTGGCTCAGATTGAACGCTGGCGGCAAGCCTAACACATGCAAGTCGAGCGGCAGCGAGAAGAGAGCTTGCTCTCTTTGTCGGCAAGCG
GCAAACGGGTGAGTAATATCTGGGGATCTACCCAAAAGAGGGGGATAACTACTAGAAATGGTAGCTAATACCGCATAATGTTGAAAAACCAAAGTGGGGG
ACCTTTTGGCCTCATGCTTTTGGATGAACCCAGACGAGATTAGCTTGTTGGTAGAGTAATAGCCTACCAAGGCAACGATCTCTAGCTGGTCTGAGAGGAT
AACCAGCCACACTGGAACTGAGACACGGTCCAGACTCCTACGGGAGGCAGCAGTGGGGAATATTGCACAATGGGCGAAAGCCTGATGCAGCTATGCCG
CGTGTATGAAGAAGGCCTTAGGGTTGTAAAGTACTTTCAGCGGGGAGGAAAAAAATAAAACTAATAATTTTATTTCGTGACGTTACCCGCAGAAGGAGCAC
CGGCTAACTCCGTGCCAGCAGCCGCGGTAATACGGAGGGTGCAAGCGTTAATCAGAATTACTGGGCGTAAAGAGCGCGTAGGTGGTTTTTTAAGTCAGG
TGTGAAATCCCTAGGCTCAACCTAGGAACTGCATTTGAAACTGGAAAACTAGAGTTTCGTAGAGGGAGGTAGAATTCTAGGTGTAGCGGTGAAATGCGTA
GATATCTGGAGGAATACCCGTGGCGAAAGCGGCCTCCTAAACGAAAACTGACACTGAGGCGCGAAAGCGTGGGGAGCAAACAGGATTAGATACCCTGG
TAGTCCATGCCGTAAACGATGTCGACTTGGAGGTTGTTTCCAAGAGAAGTGACTTCCGAAGCTAACGCATTAAGTCGACCGCCTGGGGGAGTACGGCCG
CAAGGCTAAAACTCAAATGAATTGACGGGGGCCCGCACAAGC

**> Seq18 [organism= *Buchnera aphidicola*] 16S ribosomal RNA gene [host=*Acyrthosiphon pisum* isolate Q1]**
AGAGTTTGATCCTGGCTCAGATTGAACGCTGGCGGCAAGCCTAACACATGCAAGTCGAGCGGCAGCGAGAAGAGAGCTTGCTCTCTTTGTCGGCAAGCG
GCAAACGGGTGAGTAATATCTGGGGATCTACCCAAAAGAGGGGGATAACTACTAGAAATGGTAGCTAATACCGCATAATGTTGAAAAACCAAAGTGGGGG
ACCTTTTGGCCTCATGCTTTTGGATGAACCCAGACGAGATTAGCTTGTTGGTAGAGTAATAGCCTACCAAGGCAACGATCTCTAGCTGGTCTCAGAGGATA
ACCAGCCACACTGGAACTGAGACACGGTCCAGACTCCTACGGGAGGCAGCAGTGGGGAATATTGCACAATGGGCGAAAGCCTGATGCAGCTATGCCGC
GTGTATGAAGAAGGCCTTAGGGTTGTAAAGTACTTTCAGCGGGGAGGAAAAAAATAAAACTAATAATTTTATTTCGTGACGTTACCCGCAGAAGAAGCACC
GGCTAACTCCGTGCCAGCAGCCGCGGTAATACGGAGGGTGCAAGCGTTAATCAGAATTACTGGGCGTAAAGAGCGCGTAGGTGGTTTTTTAAGTCAGGT
GTGAAATCCCTAGGCTCAACCTAGGAACTGCACTTGAAACTGGAAAACTAGAGTTTCGTAGAGGGAGGTAGAATTCTAGGTGTAGCGGTGAAATGCGTAG
ATATCTGGAGGAATACCCGTGGCGAAAGCGGCCTCCTAAACGAAAACTGACACTGAGGCGCGAAAGCGTGGGGAGCAAACAG

**> Seq19 [organism= Candidatus *Serratia symbiotica*] 16S ribosomal RNA gene [host=*Acyrthosiphon pisum* isolate Q1]**
AGAGTTTGATCCTGGCTCAGATTGAACGCTGGCGGCAGGCCTAACACATGCAAGTCGAGCGGTAGCACAAGAGAGCTTGCTCTCTGGGTGACGAGCGG
CGGACGGGTGAGTAATGTCTGGGAAACTGCCTGATGGCGGGGGATAACTAGTGGAAACGGTAGCTAATACCGCGTAACGTCGCAAGACCAAAGTGGGG
GACCTTCGGGCCTCACGCCATCAGATGTGCCCAGGTGGGATTAGCTGGTAGGTGGGGTAACGGCTCACCTAGGCGACGATCCCTAGCTGGTCTGAGAG
GATGACCAGCCACACTGGAACTGAGACACGGTCCAGACTCCTACGGGAGGCAGCAGTGGGGAATATTGCACAATGGGCGCAAGCCTGATGCAGCCATG
CCGCGTGTGTGAAGAAGGCCTTCGGGTTGTAAAGCACTTTCAGCGAGGAGAAAGGGTAATGTGTTAATAAGACATTGCATTGACGTTACTCGCAGAAGAA
GCACCGGCTAACTCCGTGCCAGCAGCCGCGGTAATACGGAGGGTGCAAGCGTTAATCGGAATTACTGGGCGTAAAGCGCACGCAGGCGGTTTGTTAAG
TCAGATGTGAAATCCCCGCGCTCAACGTGGGAACGGCATTTGAGACTGGCAAGCTAGAGTCTTGCAGAGGGGGGTAGAATTCCAGGTGTAGCGGTGAAA
TGCGTAGAGATCTGGAGGAATACCGGTGGCGAAGGCGGCCCCCAGGACAAAGACTGACGCTCAGGTGCGAAAGCGTGGGGAGCAAACAGGATTAGATA
CCCTGGTAGTCCACGCTGTAAACGATGTCGATTTGGAGGTTGCGCCCTTGAGGGGTGGCTTCCGTAGCTAACGCGTTAAATCGACCGCCTGGGGAGTAC
GGCCGcAAGGTTAAAACTCAAATGAATTGACGGGGGCCCGCACAAGCGGTGGAGCATGTGGTTTAATTCGATGCAACGCG

**> Seq20 [organism= Candidatus *Serratia symbiotica*] 16S ribosomal RNA gene [host=*Acyrthosiphon pisum* isolate Q1]**
AGAGTTTGATCCTGGCTCAGATTGAACGCTGGCGGCAGGCCTAACACATGCAAGTCGAGCGGTAGCACAAGAGAGCTTGCTCTCTGGGTGACGAGCGG
CGGACGGGTGAGTAATGTCTGGGAAACTGCCTGATGGCGGGGGATAACTAGTGGAAACGGTAGCTAATACCGCATAACGTCGCAAGACCAAAGTGGGG
GACCTTCGGGCCTCACGCCATCAGATGTGCCCAGGTAGGATTAGCTGGTAGGTGGGGTAACGGCTCACCTAGGCGACGATCCCTAGCTGGTCTGAGAG
GATGACCAGCCACACTGGAACTGAGACACGGTCCAGACTCCTACGGGAGGCAGCAGTGGGGAATATTGCACAATGGGCGCGAGCCTGATGCAGCCATG
CCGCGTGTGTGAAGAAGGCCTTCGGGTTGTAAAGCACTTTCAGCGAGGAGAAAGGGTAATGTGTTAATAAGACATTGCATTGACGTTACTCGCAGAAGAA
GCACCGGCTAGCTCCGTGCCAGCAGCCGCGGTAATACGGAGGGTGCAAGCGTTAATCGGAATTACTGGGCGTAAAGCGCACGCAGGCGGTTTGTTAAG
TCAGATGTGAAATCCCCGCGCTCAACGTGGGAACGGCATTTGAGACTGGCAAGCTAGAGTCTTGTAGAGGGGGGTAGAATTCCAGGTGTAGCGGTGAAA
TGCGTAGAGATCTGGAGGAATACCGGTGGCGAAGGCGGCCCCCTGGACAAAGACTGACGCTCAGGTGCGAAAGCGTGGGGAGCAAACAGGATTAGATA
CCCTGGTAGTCCACGCTGTAAACGATGTCGATTTGGAGGTTGCGCCCTTGAGGGGTGGCTTCCGTAGCTAACGCGTTAAATCGACCGCC

**> Seq21 [organism= Candidatus *Serratia symbiotica*] 16S ribosomal RNA gene [host=*Acyrthosiphon pisum* isolate Q1]**
AGAGTTTGATCCTGGCTCAGATTGAACGCTGGCGGCAGGCCTAACACATGCAAGTCGAGCGGTAGCACAAGAGAGCTTGCTCTCTGGGTGACGAGCGG
CGGACGGGTGAGTAATGTCTGGGAAACTGCCTGATGGCGGGGGATAACTAGTGGAAACGGTAGCTAATACCGCATAACATCGCAAGACCAAAGTGGGG
GACCTTCGGGCCTCACGCCATCAGATGTGCCCAGGTAGGATTAGCTGGTAGGTGGGGTAACGGCTCACCTAGGCGACGATCCCTAGCTGGTCTGAGAG
GATGACCAGCCACACTGGAACTGAGACACGGTCCAGACTCCTACGGGAGGCAGCAGTGGGGAATATTGCACAATGGGCGCAAGCCTGATGCAGCCATG
CCGCGTGTGTGAAGAAGGCCTTCGGGTTGTAAAGCACTTTCAGCGAGGAGAAAGGGTAATGTGTTAATAAGACATTGCATTGACGTTACTCGCAGAAGAA
GCACCGGCTAACTCCGTGCCAGCAGCCGCGGTAATACGGAGGGTGCAAGCGTTAATCGGAATTACTGGGCGTAAAGCGCACGCAGGCGGTTTGTTAAG
TCAGATGTGAAATCCCCGCGCTCAACGTGGGAACGGCATTTGAGACTGGCAAGCTAGAGTCTTGTAGAGGGGGGTAGAATTCCAGGTGTAGCGGTGAAA
TGCGTAGAGATCTGGAGGAATACCGGTGGCGAAGGCGGCCCCCTGGACAAAGACTGACGCTCAGGTGCGAAAGCGTGGGGAGCAAACAGGATTAGATA
CCCTGGTAGTCCACGCTGTAAACGATGTCGATTTGGAGGTTGCGCCCTTGAGGGGTGGCTTCCGTAGCTAACGCGTTAAATCGACCGCCTGGGGAGTAC
GGCCGCAAGGTTAAAACTCAAATGAATTGACGGGGGCCCGCACAAGC

**> Seq22 [organism= Candidatus *Serratia symbiotica*] 16S ribosomal RNA gene [host=*Acyrthosiphon pisum* isolate Q1]**
AGAGTTTGATCCTGGCTCAGATTgaACGCTGGCGGCAGGCCTAACACATGCAAGTCGAGCGGTAGCACAAGAGAGCTTGCTCTCTGGGTGACGAGCGGC
GGACGGGTGAGTAATGTCTGGGAAACTGCCTGATGGCGGGGGATAACTAGTGGAAACGGTAGCTAATACCGCATAACGTCGCAAGACCAAAGTGGGGG
ACCTTCGGGCCTCACGCCATCAGATGTGCCCAGGTGGGATTAGCTGGTAGGTGGGGTAACGGCTCACCTAGGCGACGATCCCTAGCTGGTCTGAGAGG
ATGACCAGCCACACTGGAACTGAGACACGGTCCAGACTCCTACGGGAGGCAGCAGTGGGGAATATTGCACAATGGGCGCAAGCCTGATGCAGCCATGC
CGCGTGTGTGAAGAAGGCCTTCGGGTTGTAAAGCACTTACAGCGAGAAGAAAGGGTAATGTGTTAATAAGACATTGCATTGACGTTACTCGCAGAAGAAG
CACCGGCTAACTCCGTGCCAGCAGCCGCGGTAATACGGAGGGTGCAAGCGTTAATCGGAATTACTGGGCGTAAAGCGCACGCAGGCGGTTTGTTAAGT
CAGATGTGAAATCCCCGCGCTCAACGTAGGAACGGCATTTGAGACTGGCAAGCTAGAGTCTTGTAGAGGGGGGTAGAATTCCAGGCGTAGCGGTGAAAT
GCGTAGAGATCTGGAGGAATACCGGTGGCGAAGGCGGCCCCCTGGACAAAGACTGACGCTCAGGTGCGAAAGCGTGGGGAGCAAACAGGATTAGATAC
CCTGGTAGTCCACGCTGTAAACGATGTCGATTTGGAGGTTGCGCCCTTGAGGGGTGGCTTCCGTAGCTAACGCGTTAAATCGACCGCCTGGGGAGTACG
GCCGCAAGGTTAAAACTCAAATGAATTGACGGGGGCCCGCACAAGCGGTGGaGCATGTGGTTTAATTCGATGCAACGCGA

**> Seq23 [organism= Candidatus *Serratia symbiotica*] 16S ribosomal RNA gene [host=*Acyrthosiphon pisum* isolate Q1]**

GGCCTAACACATGCAAGTCGAGCGGTAGCACAAGAGAGCTTGCTCTCTGGGTGACGAGCGGCGGACGGGTGAGTAATGTCTGGGAAACTGCCTGATGG
CGGGGGATAACTAGTGGAAACGGTAGCTAATACCGCATAACGTCGCAAGACCAAAGTGGGGGACCTTCGGGCCTCACGCCATCAGATGTGCCCAGGTA
GGATTAGCTGGTAGGTGGGGTAACGGCTCACCTAGGCGACGATCCCTAGCTGGTCTGAGAGGATGACCAGCCACACTGGAACTGAGACACGGTCCAGA
CTCCTACGGGAGGCAGCAGCGGGGAATATTGCACAATGGGCGCAAGCCTGATGCAGCCATGCCGCGTGTGTGAAGAAGGCCTTCGGGTTGTAAAGCAC
CTTCAGCGAGGAGAAAGGGTAATGTGTTAATAAGACATTGCATTGACGTTACTCGCAGAAGAAGCACCGGCTAACTCCGTGCCAGCAGCCGCGGTAATA
CGGAGGGTGCAAGCGTTAATCGGAATTACTGGGCGTAAAGCGCACGCAGGCGGTTTGTTAAGTCAGATGTGAAATCCCCGCGCTCAACGGGGGAACGG
CATTTGAGACTGGCAAGCTAGAGTCTTGTAGAGGGGGGTAGAATTCCAGGTGTAGCGGTGAAATGCGTAGAGATCTGGAGGAATACCGGTGGCGAAGG
CGGCCCCCTGGACAAAGACTGACGCTCAGGTGCGAAAGCGTGGGGAGCAAACAGGATTAGATACCCTGGTAGTCCACGCTGTAAACGATGTCGATTTGG
AGGTTGCGCCCTTGAGGGGTGGCTTCCGTAGCTAACGCGTTAAATCGACCGCCTGGGGAGTACGGCCGCAAGGTTAAAACTCAAATGAATTGACGGGG
GCCCGCACAAGCGGTGGAGCATGT

**Table S1. Summary of the statistical model of aphid immunity**. Analysis of Deviance Table (Type II Wald chi-square tests) of the mixed effect generalised linear model (Model1) investigating the effects of intraguild predation by the aphid lion (No [without IGP], Yes [with IGP]), pea aphid lineage (N116, Q1), parasitoid genotype and the two-way interactions between the main effects on aphid immunity ratio (IR). Resulting from a full quantitative genetic mating design, with n sires being mated to at least three dams, the sire × dam mating groupings produced the intraspecific genetic variability in the parasitoid (genotype/daughters [sibs, and half-sibs]). IR in the microcosm is the proportion of healthy aphids (non-mummified *i.e.* unparasitoidised) after 11 days of exposure to the parasitoid genotype relative to the entire population of aphids (healthy and mummified) per aphid lineage. Significant results are shown in bold.

| | $\chi^2$ | Df | Pr(>Chisq) |
|---|---|---|---|
| IGP | 1.74 | 1 | 0.188 |
| Aphid Lineage | 28.5 | 1 | **< 0.0001** |
| Parasitoid Genotype | 95.76 | 37 | **< 0.0001** |
| Aphid Lineage x Parasitoid Genotype | 12.1 | 3 | **0.007** |
| IGP x Parasitoid Genotype | 37.23 | 15 | **0.001** |
| IGP x Aphid Lineage | 2.03 | 1 | 0.155 |

**Table S2. Summary of the statistical model of aphid mummy position.** Analysis of Deviance Table (Type II tests) of the generalised linear model (Model2, specified in the main text) applied to investigate the effects of intraguild predation by aphid lion (No [without IGP], Yes [with IGP]), pea aphid lineage (N116, Q1), parasitoid genotype and the presented interactions between the main effects on the proportions of mummified aphid off the host plant at the 11$^{th}$ day of the experiment. Resulting from a full quantitative genetic mating design, with n sires being mated to at least three dams, the sire × dam mating groupings produced the intraspecific genetic variability in the parasitoid (genotype/daughters [sibs, and half-sibs]). Significant results are shown in bold, marginally significant results are shown in italics.

| | $LR_{\chi^2}$ | Df | Pr(>Chisq) |
|---|---|---|---|
| IGP | 0.29 | 1 | 0.59 |
| Aphid Lineage | 7.05 | 1 | **0.008** |
| Parasitoid Genotype | 40.62 | 28 | *0.058* |
| Aphid Lineage x Parasitoid Genotype | 4.373 | 2 | 0.112 |
